# Allele-specific collateral and fitness effects determine the dynamics of fluoroquinolone-resistance evolution

**DOI:** 10.1101/2020.10.19.345058

**Authors:** Apostolos Liakopoulos, Linda B. S. Aulin, Matteo Buffoni, Efthymia Fragkiskou, J. G. Coen van Hasselt, Daniel E. Rozen

**Affiliations:** Department of Microbial Biotechnology and Health, Institute of Biology, Leiden University, Leiden, The Netherlands; Leiden Academic Centre for Drug Research, Leiden University, Leiden, The Netherlands

**Keywords:** antibiotic resistance, collateral sensitivity, population dynamics, pharmacokinetics/pharmacodynamics, *Streptococcus pneumoniae*

## Abstract

Collateral sensitivity (CS), which arises when resistance to one antibiotic increases sensitivity towards other antibiotics, offers novel treatment opportunities to constrain or reverse the evolution of antibiotic-resistance. The applicability of CS-informed treatments remains uncertain, in part because we lack an understanding of the generality of CS effects for different resistance mutations, singly or in combination. Here we address this issue in the Gram-positive pathogen *Streptococcus pneumoniae* by measuring collateral and fitness effects of clinically relevant *gyrA* and *parC* alleles, and their combinations, that confer resistance to fluoroquinolones. We integrated these results in a mathematical model which allowed us to evaluate how different *in silico* combination treatments impact the dynamics of resistance evolution. We identified common and conserved CS effects of different *gyrA* and *parC* alleles; however, the spectrum of collateral effects was unique for each allele or allelic pair. This indicated that allelic identity can impact the evolutionary dynamics of resistance evolution during monotreatment and combination treatment. Our model simulations, which included the experimentally derived antibiotic susceptibilities and fitness effects, and antibiotic specific pharmacodynamics, revealed that both collateral and fitness effects impact the population dynamics of resistance evolution. Overall, we provide evidence that allelic identity and interactions can have a pronounced impact on collateral effects to different antibiotics and suggest that these need to be considered in models examining CS-based therapies.

**Significance:** A promising strategy to overcome the evolution of antibiotic resistant bacteria is to use collateral sensitivity-informed antibiotic treatments that rely on cycling or mixing of antibiotics, such that that resistance towards one antibiotic confers increased sensitivity to the other. Here, focusing on multi-step fluoroquinolone resistance in *Streptococcus pneumoniae*, we show that antibiotic-resistance induces diverse collateral responses whose magnitude and direction are determined by allelic identity. Using mathematical simulations, we show that these effects can be exploited via combination treatment regimens to suppress the *de novo* emergence of resistance during treatment.

## Introduction

Antibiotics are a cornerstone of the prevention and treatment of bacterial infections; however, their efficacy is rapidly declining due to the emergence and spread of antibiotic resistance (1). Advances in antibiotic discovery and design have not kept pace with resistance evolution (2), necessitating new experimentally validated treatment strategies (3). Selection inversion, a strategy that aims to circumvent the emergence and dissemination of antibiotic resistance by combining existing antibiotics based on their physiological and/or evolutionary interactions, has recently gained prominence (4). Among these, collateral sensitivity-informed strategies are particularly promising (5).

Collateral sensitivity (CS) occurs when mutations conferring resistance to one antibiotic increase sensitivity towards other antibiotics in the same or different functional class (6). Because of this effective trade-off, bacteria treated with a pair of drugs exhibiting reciprocal CS are unable to simultaneously evolve resistance to both agents (4). Recent studies have examined the frequency and mechanisms of CS, although these have been largely restricted to laboratory strains of a small number of bacterial species (5, 7–14). In addition, none of these studies have examined if collateral effects are conserved for different clinically circulating resistance alleles to the same antibiotics. This is especially relevant given that resistance to any given antibiotic can arise from mutations in different genes or at different sites within a gene (15). Because these resistance alleles cause distinct phenotypic effects, both with respect to changes in MIC and bacterial fitness (16), they may also underlie a distinct spectrum of collateral effects, only some of which would be suitable for CS-informed therapies. To address this limitation, we examine collateral responses to distinct genes and alleles that confer resistance to fluoroquinolones (FQ) in the Gram-positive pathogen *Streptococcus pneumoniae*.

*S. pneumoniae* invasive infections are responsible for the most deaths among vaccine-preventable diseases globally (17). Fluoroquinolones are a mainstay of treatment against invasive pneumococcal disease (18), but successful FQ treatment is threatened by the emergence of FQ-resistant strains, which have been reported to be as high as 10.5% (19). *De novo* FQ-resistance in *S. pneumoniae* arises predominantly via the stepwise accumulation of chromosomal mutations in the quinolone resistance-determining regions (QRDRs) of the DNA gyrase (*gyrA*) and/or topoisomerase IV (*parC*) genes (19). Mutations in either *gyrA* or *parC* result in low-level FQ-resistance, whereas the combination of mutations in both genes results in high-level resistance, often affecting multiple agents within the class (18, 20). Alleles carrying different mutations in QRDR regions, in pneumococci and other species, have varying impacts on FQ-resistance and on the fitness of strains carrying them (21, 22), effects that are believed to influence the population frequencies of different alleles during monotherapy (23). If similar phenotypic heterogeneity exists for collateral responses, this could impact the efficacy of CS-based combination therapies and the persistence and spread of FQ-resistance.

Here, we studied collateral effects of FQ-resistance by generating *S. pneumoniae* mutant strains via allelic replacement that encode clinically relevant FQ-resistance mutations in *gyrA* and *parC* (19). These mutants were used to assess how different alleles conferring resistance to the same antibiotic influence the susceptibility to other antibiotics and whether interactions between these alleles modulate their collateral effects. We then developed and applied a mathematical model to quantitatively study the population dynamics of resistance evolution during different antibiotic combination treatment regimes.

## Results

### Extensive and conserved collateral effects to fluoroquinolone-resistance

To investigate collateral effects to FQ resistance in *S. pneumoniae*, we generated a panel of 16 D39 strains harboring mutations in the QRDRs of *gyrA* and/or *parC* by natural transformation (**Figure 1A**). Specifically, we generated eight strains encoding alleles with FQ-resistance associated mutations in either gyrA (gx, with x = S81F, S81Y, E85G or E85K) or parC (py, with y = S79F, S79Y, D83N or D83Y) and eight strains with combinations of both alleles (gxpy). Each strain is hereafter denoted based on its FQ-resistance related amino acid mutations. All mutant strains had decreased susceptibility to ciprofloxacin, where the gxpy showed the most pronounced MIC increase (32 - 64 mg/L), followed by py (16 - 32 mg/L), and lastly gx (MIC 8 mg/L) (**Figure 1B**). In addition, we observed significant fitness reductions in transformed strains (**Figure 1C**), with gxpy strains exhibiting the lowest mean relative growth rate (0.56; with a range between 0.29 - 0.74), followed by gx (0.60; 0.41 - 0.77) and lastly py (0.85; 0.39 - 1.47) strains.

**Figure 1.**
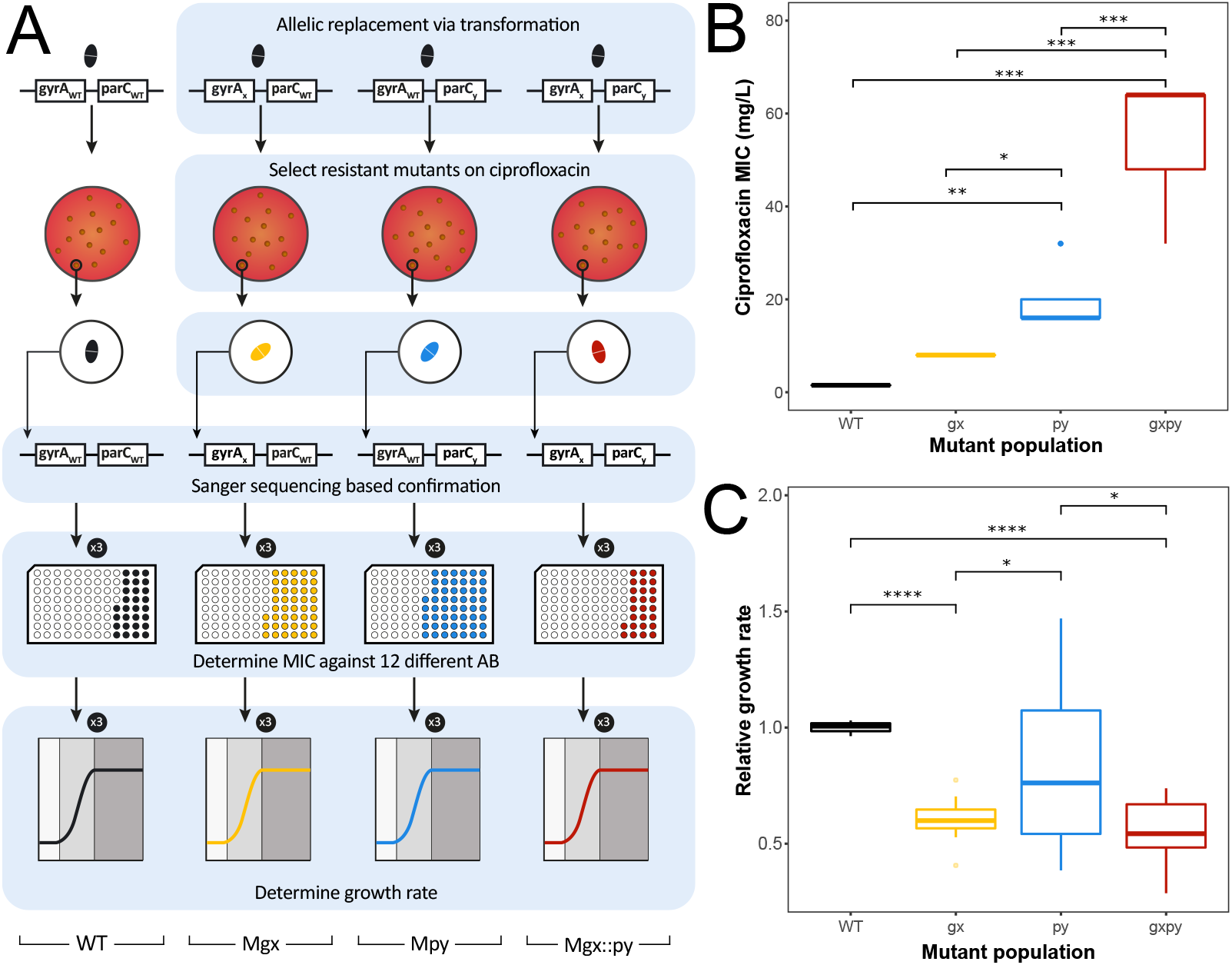
Experimental design and determined MIC and growth rates. **A:** Overview of experimental workflow. Mutants of *S. pneumoniae* D39 strain encoding *gyrA* (gx, blue), *parC* (py, orange) or double *gyrA*::*parC* (gxpy, red) alleles that confer FQ-resistance were generated via transformation followed by allelic replacement. Mutants were subjected to antimicrobial susceptibility testing for 12 clinically relevant antibiotics and ciprofloxacin. A decrease in MIC of each mutant relatively to that of the wild-type D39 strain (WT, green) was determined as collateral sensitivity (CS) and an increase as collateral resistance (CR). The growth rate of the WT and each mutant was determined to estimate *in vitro* fitness. **B:** Ciprofloxacin MICs for the wild-type *S. pneumoniae* D39 (WT, black), gyrA mutants (gx, yellow), parC mutants (py, light blue), and their corresponding gyrA::parC double-allele mutant (gxpy, red). **C:** Relative growth rate of *gyrA* (gx, yellow), *parC* (py, light blue), and *gyrA*::*parC* (gxpy, red) mutants compared to WT (black) using a two-sided t-test. Significant differences are indicated by * (p<0.05), ** (p<0.01), *** (p<0.001) and **** (p <0.0001).

The collateral effects of these mutants were assessed against 12 antibiotics belonging to a broad range of classes, including commonly used anti-pneumococcal agents. Collateral effects were observed in approximately 52% of the possible instances (100 out of 192) against all tested antibiotics (**Figure 2**). Among these, 87% were CS and the remaining 13% CR. For 7 of 12 antibiotics, collateral effects were in the same direction for all mutants (CS), whereas for the remaining antibiotics collateral effects were mixed between CS and CR. All mutants (100%) exhibited CS to gentamicin, 81.3% to clindamycin, 75.0% to tetracycline, 56.3% to trimethoprim/sulfamethoxazole, 43.8% to penicillin and fusidic acid, and only 6.3% to vancomycin.

**Figure 2.**
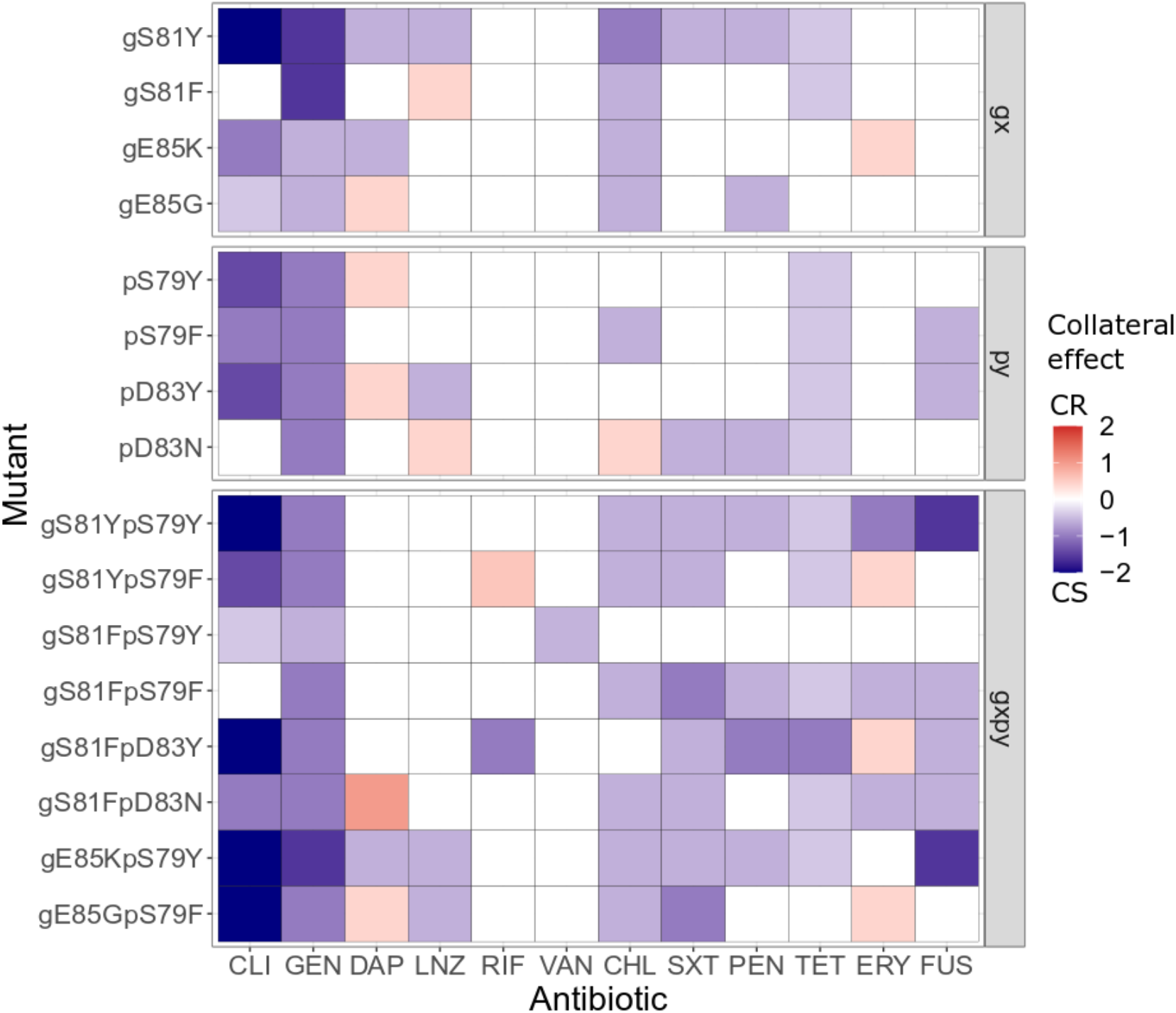
Collateral effect (CE) profiles for FQ-resistant *S. pneumoniae,* including *gyrA* (gx) and *parC* (py) mutants, and *gyrA*::*parC* double mutants (gxpy). Antibiotics are ranked based on the hierarchical clustering of their associated collateral effects. Color indicates the magnitude of the collateral effect quantified as the mean log2 relative change of MIC compared to the WT. Red indicates collateral resistance (CR) and blue collateral sensitivity (CS).

Single-allele strains, gx and py, exhibited CS to a median of four antibiotics with log-scale fold changes relative to the isogenic WT varying from -0.42 to -1.58, and CR to a median of one antibiotic with fold change of 0.42. Double gxpy strains exhibited CS to a median of seven antibiotics with log-scale fold changes relative to the isogenic WT varying from -0.42 to -2 and CR to a median of 0.5 antibiotic with fold changes of 0.42 to 1. Conserved CS effects to gentamicin and clindamycin were pronounced, with a median log-fold change in MICs of -1.2 and -1 respectively.

Hierarchical clustering revealed four antibiotic groups for which resistant mutants exhibited similar collateral effects (**Figure S1**); only one of these groups consisted exclusively of functionally similar antibiotics, including the protein synthesis inhibiting antibiotics, gentamicin and clindamycin, confirming that collateral effects cannot be simply predicted based on the antibiotic target. Single- and double-allele mutants did not cluster distinctively based on their effects within the five strain clusters that we observed (**Figure S1**); this suggests that many of the collateral effects in double mutants are potentially caused by the mutations of each single allele, although the magnitude of these effects is altered in the double-mutants, as discussed below.

### Heterogeneity in collateral effects to fluoroquinolone-resistance

Despite the extensive and conserved collateral effects that we observed, we found that the single-allele gyrA and parC strains exhibited heterogeneity in their collateral effects even between strains carrying different resistant alleles of the same gene (**Figure 2**). For instance, strains carrying the gyrA alleles S81F and S81Y exhibited CS respectively to three and eight antibiotics and a median fold decrease of -0.58 in MICs. Similarly, the strain carrying the parC allele S79Y exhibited CS to three antibiotics with a median fold decrease of -1 in MICs, whereas the strain carrying the S79F allele showed CS to five antibiotics with a median fold decrease of -0.58. Interestingly, the heterogeneity was not limited to the number or intensity but also the direction of the effects. For example, strains carrying the gyrA S81F and S81Y alleles exhibited CR and CS to linezolid, respectively; a similar difference was observed with daptomycin for strains carrying the gyrA E85G and E85K alleles.

Collateral effects among double-allele gxpy strains (**Figure 2**) varied in their number, intensity and/or direction as well. For instance, the double-allele strain gS81FpS79F exhibited CS respectively to seven antibiotics with a median fold decrease of -0.58 in MICs whereas the strain gS81FpS79Y showed CS to three antibiotics with a median fold decrease of -0.57. Heterogeneity in the direction of the collateral effects was observed for erythromycin where several of the double-allele strains exhibited either CS (gS81YpS79Y, gS81FpS79F, gS81FpD83N) or CR (gS81YpS79F, gS81FpD83Y, gE85GpS79F). In addition, collateral effects in double-allele strains appeared to differ from those expected based on the strains carrying the respective single alleles, resulting in either lower (*i.e.,* gS81YpS79Y for fusidic acid) or higher (*i.e.,* gS81YpS79F for gentamycin) MIC for the collaterally affected antibiotic.

### Antibiotic-resistance linked fitness does not predict collateral effects

All gyrA and parC alleles, aside from pS79F, caused significantly reduced growth rates compared to the WT strain (**Figures 1C and S2**). In addition, significant differences (ANOVA, p<0.05) were observed between strains carrying different FQ-resistant alleles of either the *gyrA* (gS81F and gE85K |ΔK_G_| = 0.0134 h^-1^) or the *parC* (pS79F and pD83N |ΔK_G_| = 0.0663 h^-1^, pS79F and pD83Y |ΔK_G_| = 0.0431 h^-1^, pS79Y and pS79F |ΔK_G_| = 0.0461 h^-1^), but not between the strains carrying resistant alleles of both genes (**Figure S2**). Our analysis indicated that there was no clear overall correlation between the fitness of the generated strains and their collateral effects (**Figure S3**). This suggests that the fitness costs of resistance have only a limited impact on observed CS effects overall, with this impact being heterogeneous and antibiotic-dependent.

### Combination treatment with clinical dosing regimens suppresses *de novo* antibiotic-resistance development

Given the distinct collateral and fitness effects of FQ resistance alleles, we used simulations to assess the emergence and fixation of resistant strains based on eight evolutionary trajectories leading to high-level FQ-resistance (**Figure 3A**), where each evolutionary trajectory included one double-allele mutant and its corresponding single-allele mutants. To this end, we developed a mathematical model and studied the impact of these differences on treatment outcomes (**Figure 3B**) under a clinical dosing schedule for ciprofloxacin monotreatment (500 mg b.i.d., Css 1.39 mg/L), and in combination with erythromycin (600 mg b.i.d., Css 0.48 mg/L), linezolid (600 mg b.i.d, Css 7.33 mg/L), and penicillin (3000 mg b.i.d., Css 6.95 mg/L). The model incorporated our experimentally derived antibiotic MICs, collateral and fitness effects, and antibiotic-specific pharmacodynamic relationships (details and estimated parameters can be found in Appendixes 1 and 2). For the different simulation scenarios, we then computed the probability of resistance evolution for each of the eight evolutionary trajectories leading to high-level FQ-resistance (ET) shown in **Figure 3C**. Resistance was defined as the end of treatment fixation of the gx, py and/or gxpy subpopulation, which occurred when the subpopulation-specific bacterial density exceeded the resistance cut-off. The resistance cut-off was set to 10^4^ cfu/mL and represents an established infection (24).

**Figure 3.**
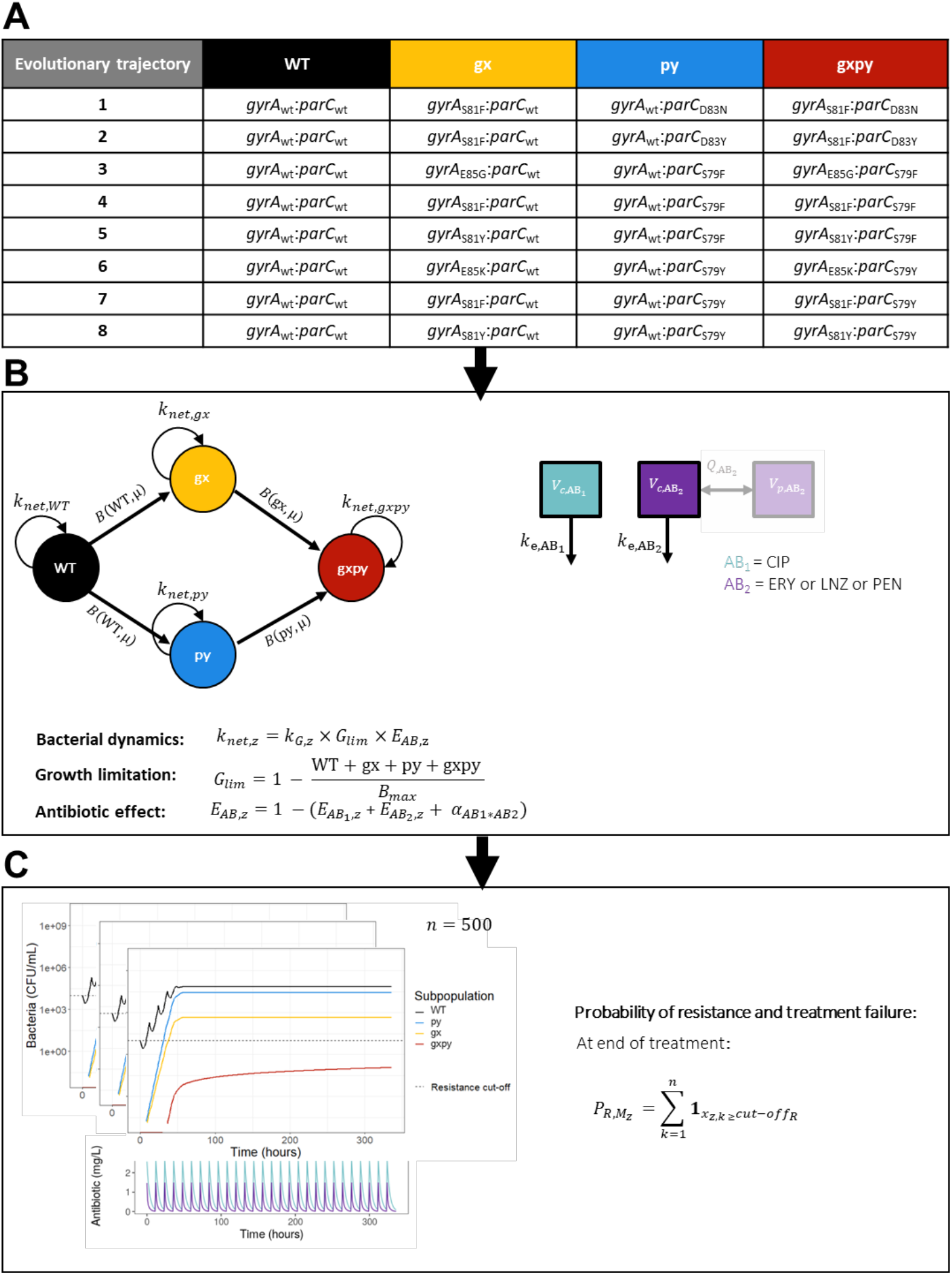
Overview of treatment simulation workflow. **A**: Trajectories leading to high-level FQ-resistance (ETs) used for simulations, with each trajectory consisting of four subpopulations representing the wild-type *S. pneumoniae* D39 (WT, green), a *gyrA* mutant (gx, yellow), a *parC* mutant (py, light blue), and their corresponding *gyrA*::*parC* double-allele mutant (gxpy, red). The experimentally determined MICs and relative growth rate (KG) for each specific strain were used to inform the model. **B:** Schematic of the stochastic hybrid ordinary deferential equation (ODE) model use for simulations. The bacterial model comprised of ODEs describing the change of bacterial density over time for each subpopulation. Bacterial growth was modeled using a capacity limitation (*Glim*) and a subpopulation specific growth rate (*kG,z*). Unidirectional mutations were implemented as a stochastic event using random sampling from a binomial distribution with a probability representing the mutation frequency (*μ*) and sample size equal to the density of each bacterial subpopulation. The bacterial model was linked to a pharmacokinetic model of ciprofloxacin (CIP) and a pharmacokinetic model of erythromycin (ERY), linezolid (LNZ), or penicillin (PEN). Antibiotic effect (*EAB,z*) was included on each subpopulation and was driven by the simulated plasma concentrations. Pharmacodynamic drug interactions were included for drug combinations for which significant non-additivity was identified. **C:** Treatments were simulated 500 times per evolutionary trajectory. Pharmacokinetic profiles of an example combination treatment are depicted in the lower graph, while the n^th^ realization of the bacterial dynamics corresponding to the example treatment are shown in the top graph. The probability of resistance (mutants) or failure of eradication (WT) was calculated for each trajectory and treatment by summarizing the number of simulations for which bacterial concentrations exceeded the resistance cut-off (dashed line) at end of the treatment for each individual subpopulation.

Our simulations of clinical ciprofloxacin dosing schedules, where Css equates to 0.9-fold of WT MIC (1.5 mg/L), revealed that monotreatment failed to eradicate the WT and promoted resistance evolution (**Figure 4A-B**) for all evolutionary trajectories due to the emergence of low-level FQ-resistant mutants (gx and/or py). However, adding a second drug effectively eradicated the WT (**Figure 4C**). Treatments using linezolid (1.5 mg/L, 4.9 x WT MIC) or penicillin (0.012 mg/L, 594 x WT MIC) resulted in 0% probability of resistance, whereas erythromycin treatment (0.19 mg/L, 2.6 x WT MIC) promoted the emergence of either exclusively py (75.6-80.4%) or both py (75.6%) and gx (77%) strains for all but one evolutionary trajectory (ET 1). The emergence of gxpy double-allele strains was only observed for a single trajectory (ET 2), with a very low probability of resistance (0.2%). Overall, our findings support the idea that antibiotic combinations can minimize the risk of *de novo* antibiotic-resistance evolution, but the benefit depends on the specific antibiotic combination.

**Figure 4.**
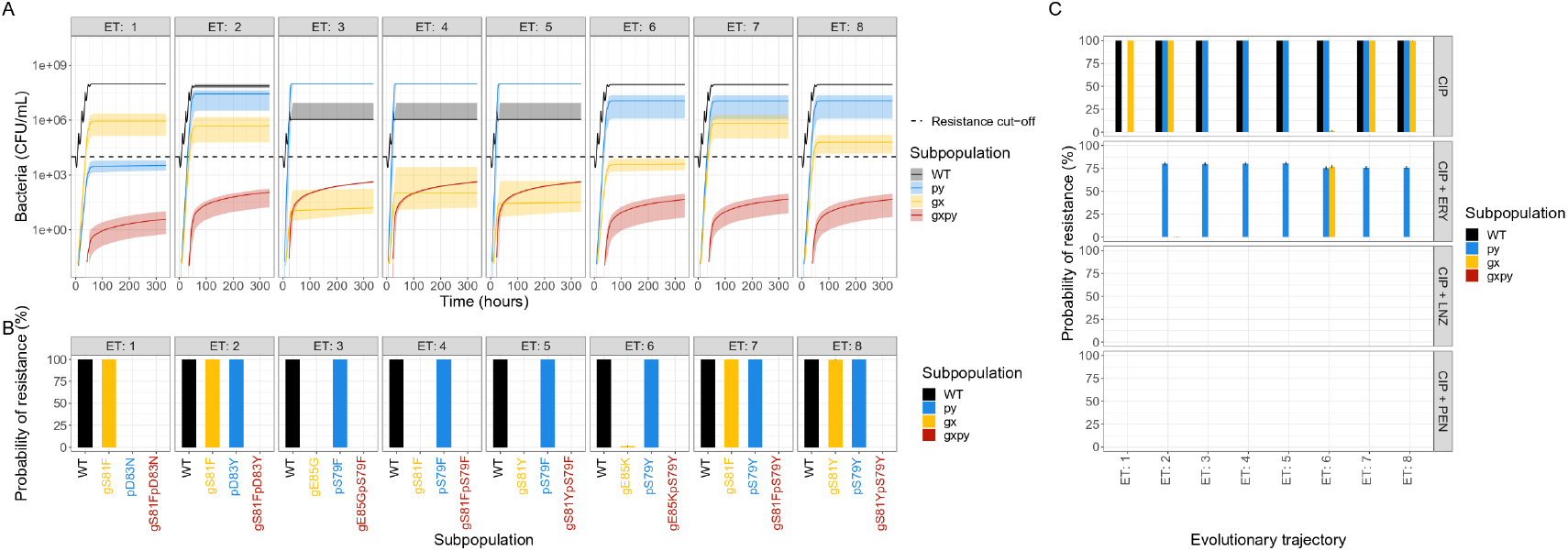
Treatment outcomes of simulated clinical ciprofloxacin mono- or combination therapies. Each simulation includes WT (black), a *gyrA* mutant gx (yellow), a *parC* mutant py (light blue), and the double-allele mutant gxpy (red). **A**: Treatment outcomes of simulated clinical ciprofloxacin monotreatment (CIP, 500 mg b.i.d., average unbound steady state concentrations (C_ss_) 1.39 mg/L) vary between different trajectories (ETs). Solid line represents the median bacterial growth dynamics of 500 simulations where bacterial subpopulations are indicated by the respective color. The shaded areas represent the 5^th^ and 95^th^ simulated percentiles. The black dashed line represents the resistance cut-off, which is equal to the WT bacterial density at start of treatment. Although the bacterial dynamics and resistance development differed between the ETs, the simulated ciprofloxacin treatment was ineffective for all scenarios. **B:** End-of-treatment probability of resistance (mutants) and WT eradication failure, indicated in gray, was calculated for different ETs (panels), and subpopulations (x-axis) separately. **C:** Treatment outcome for clinical dosing regimens of ciprofloxacin monotreatment or combination therapies. Simulation outcomes with clinically relevant dosing regimens of ciprofloxacin as monotreatment and in combination with erythromycin (ERY, 600 mg b.i.d., Css 0.48 mg/L), linezolid (LNZ, 600 mg b.i.d., Css 7.33 mg/L), or penicillin (PEN, 3 g b.i.d., Css 6.95 mg/L). CIP as monotreatment had a high probably of resistance, foremost selecting for strains encoding FQ-resistant *parC* alleles (py, light blue) and secondly *gyrA* alleles (gx, yellow). The addition either LNZ or PEN suppressed the probability of resistance for all evolutionary trajectories whereas that of ERY resulted in high probability of resistance (>75%) for at least one of the single-allele mutants for seven out of the eight trajectories. All treatment combinations eradicated the WT (black) and suppressed the double-allele (gxpy, red) subpopulations.

### Probability of *de novo* resistance evolution varies between trajectories

Next, to examine the effects of CS-informed treatments using lower antibiotic concentrations due to collateral effects, we simulated combination treatments with Css ranging from 0.25 to 4 x MIC (*i.e.,* -2 to 2 log2 MIC) of the WT population. Our simulations indicated that eradication of WT bacteria was primarily observed for dosing schedules with Css of at least one of the antibiotics being equal to or larger than the MIC of the WT population (**Figure 5A-B, Figure S4A-B**). At these concentrations a clear impact of collateral effects was observed. Specifically, subpopulations with CS were suppressed while subpopulations with CR were associated with a high probability of resistance evolution.

**Figure 5.**
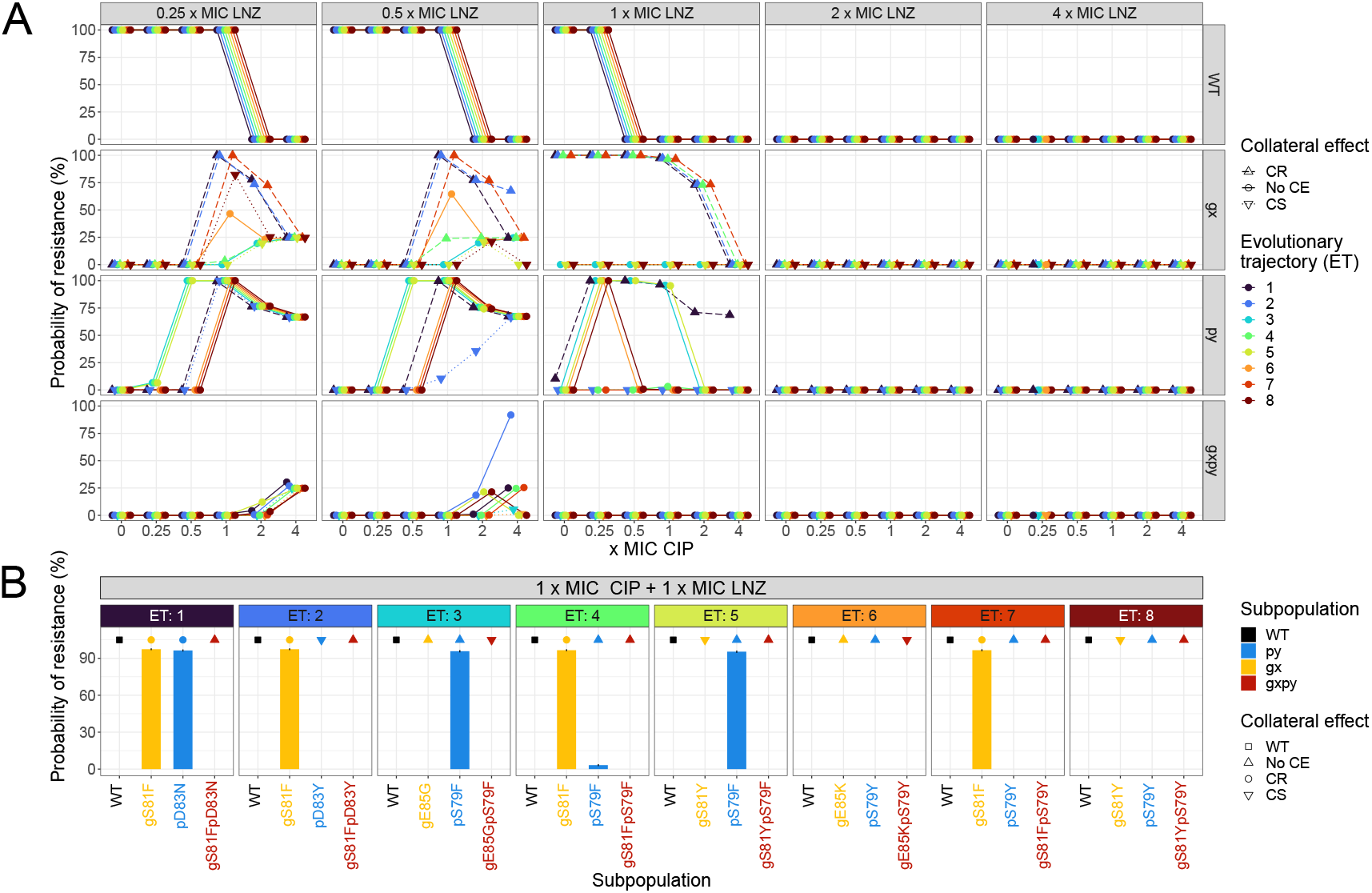
Treatment outcome of ciprofloxacin (CIP) in combination with linezolid (LNZ). The treatments were evaluated on eight different trajectories leading to high-level FQ-resistance (see Figure 4A). **A:** Simulation outcomes of CIP in combination with LNZ with average unbound steady state concentrations (C_ss_) around the MIC of the WT population (C_ss_ 0.25 - 4 x MIC). Concentration dependency related to WT strain eradication and resistance evolution. Each point represents a bacterial subpopulation from a specific evolutionary trajectory (ET, indicated by color). Collateral effects (CE) are indicated by shape and line-type, where solid lines with round points represent no CE, dashed lines with triangles pointing up collateral resistance (CR), and dotted lines with triangles pointing down collateral sensitivity (CS). **B:** End-of-treatment probability of resistance (mutants) and treatment failure (WT) for different evolutionary trajectories treated with antibiotic doses resulting in an average steady state plasma concentration of CIP and LNZ equal to the 1 x MIC of the WT. The presence or not of collateral effects (CE) are indicated by text in the box above each specific mutant. The evolutionary trajectory (ET) *i.e*., the multistep alternative pathways for the *de novo* emergence of a particular high-level FQ-resistant double-mutant (gxpy), had an impact on treatment outcome, although it was not the sole determinant.

In addition, our simulations showed that the population dynamics and treatment outcomes for ciprofloxacin mono- and combination treatments with Css around the MIC of the WT population varied both within and between antibiotic treatments and evolutionary trajectories (**Figure S5, Figure S6**). In particular, the fixation of each specific gx and py strain in the population varied between trajectories due to competition between these two arising clonal lineages. For example, when comparing our eight different evolutionary trajectories treated with 1 x MIC for both ciprofloxacin and linezolid (**Figure 5B**), the fixation of specific gx, and therefore the emergence of resistance, largely differed depending on the concurrent py strain. The specific treatment schedule resulted in high probabilities of fixation for at least one FQ-resistant subpopulation (>95%) for six out the eight evolutionary trajectories. However, the probability of fixation for different strains varied across these trajectories, *e.g.,* whereas the probability of fixation of pS794 was 0.032% when gS81F was the concurrent strain (ET 4), it rose to 98.5% when gE85G was the concurrent strain (ET 3). Overall, these findings suggest that strain-specific collateral and fitness effects as well as clonal competition have an impact on the dynamics of FQ-resistance evolution.

### Collateral and fitness effects determine the outcome of antibiotic combination treatments

The heterogeneous fixation of individual strains observed (**Figure 4** **Figure 5** **Figure S4, Figure S5**, and **Figure S6**) prompted us to assess and disentangle the impact of collateral and fitness effects on this phenomenon. To this end, we simulated four different scenarios of treatment combinations with ciprofloxacin (1 x MIC) and linezolid (1 x MIC), ciprofloxacin (1 x MIC) and erythromycin (1 x MIC), as well as ciprofloxacin (0.5 x MIC) and penicillin (1 x MIC). Specifically, the first scenario (Scenario 1) did not include the experimentally derived collateral or fitness effects, while the remaining simulations included either the fitness (Scenario 2) or the collateral effect (Scenario 3), or both (Scenario 4). Two different model structures were used for the simulations of each scenario, one simplified where each evolutionary trajectory was evaluated separately as described in **Figure 3A** allowing a low degree of clonal competition (Approach A) but high power to disentangle impact of collateral and fitness effects, and an adapted where all strains used in our study were included simultaneously (free-for-all simulation; Approach B) allowing for a higher degree of clonal competition, capturing a more real-life scenario compared to our eight-defined-trajectory approach.

When simulating each evolutionary trajectory separately and in the absence of both collateral and fitness effects (Scenario 1A), the ciprofloxacin-linezolid treatment (**Figure 6A**) resulted in a fixation probability of 47.4% for py strains for all evolutionary trajectories but ET 1, with a probability of resistance of 89.0%. Notably, ET 1 is the only trajectory including pD83N, which is the single-allele strain associated with the highest ciprofloxacin MIC (32 mg/L) in our experiments. Both the gx and gxpy strains were completely suppressed in these simulations. For the two other simulated antibiotic combinations (**Figure S7**), gx and py strains had similar probability of resistance (∼80%), whereas the gxpy strains had probability lower than 1%. The latter suggests that due to the low antibiotic pressure in our simulations by the time a double mutant emerges via a two-step mutation, the single mutant populations are large enough to suppress its fixation. The outcome of these simulations suggests that the majority of the heterogeneity observed in the previous simulations are due to the collateral and/or fitness effects, and not differences in sensitivity towards ciprofloxacin. In the free-for-all simulation (Scenario 1B), we observed similar high probability of resistance development during both ciprofloxacin-erythromycin (>80%) and ciprofloxacin-penicillin combination treatments (>70%) for all the single-allele strains tested (**Figure 6B and Figure S8**). During ciprofloxacin-linezolid combination treatment however, the highest probability of resistance was due to the fixation of the pD83N strain (89.2%), followed by that of other py strains (pS79F, pS79Y, and pD83Y) with resistance probabilities close to 50%, while the other FQ-resistant strains had a 0% probability of fixation (**Figure 6B**).

**Figure 6.**
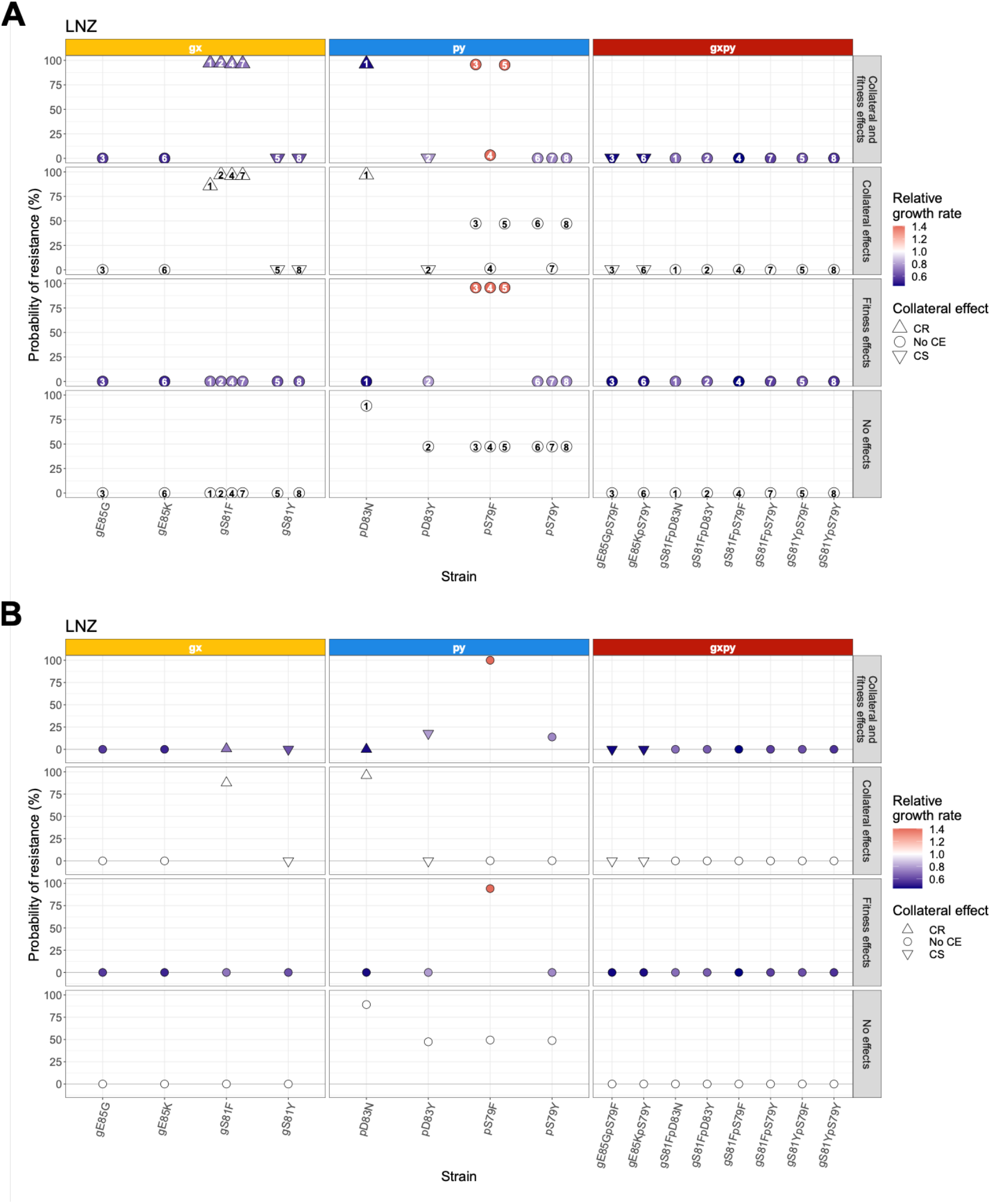
Relationship between collateral effect, relative growth rate, and probability of resistance under treatment with antibiotic combinations. **A:** Simulation of the eight defined *S. pneumoniae* evolutionary trajectories (ET) leading to high-level FQ-resistance (indicated by number). **B:** Free-for-all simulation of resistance evolution in *S. pneumoniae* allowing for the simultaneous emergence of 16 different FQ-resistant subpopulations (four gx, four py and eight gxpy). In both simulations, the treatment combinations consisted of ciprofloxacin (CIP) and linezolid (LNZ) with average steady state plasma concentrations equal to 1 x MIC of the WT. Each unique strain is represented on the x-axis. The collateral effect (CE) of LNZ is depicted using shapes, where tringle point up is collateral resistance (CR), pointing down is collateral sensitivity, and circle represents no CE. The probability of resistance at the end of treatment is shown on the y-axis. Colors indicate the relative growth rate of each mutant compared to the WT. Each panel column denotes a FQ-resistant subpopulation (gx, py or gxpy) with the top panels representing simulations where both collateral and fitness effects were included, top mid panels representing simulations based solely on the experimentally determined collateral effects, lower mid panels based solely on the experimentally determined relative growth rate, and the lowest panel simulations where neither collateral nor fitness effects were included.

Reintroducing fitness effects in the absence of collateral effects and simulating each trajectory separately (Scenario 2A) reveled that only the strain with a fitness advantage (pS79Y, ET 3, 4, and 5) fixed during the simulated ciprofloxacin-linezolid treatment (probability of resistance 95.8%) (**Figure 6A**). This differed from the other two treatments simulated for this scenario, where there was large variation between trajectories and specific strains, *e.g.*, under ciprofloxacin-penicillin treatment gS81F paired with pS79F (ET 4) was associated with a probability of resistance of 16.2%, while when paired with pD83Y (ET 2) or pS79Y (ET 7), the probability was 59.2% and 76.6% (**Figure S7**), respectively. These inter-trajectory differences relating to specific strains again highlight the importance of clonal competition in resistance evolution. During the free-for-all simulation (Scenario 2B), only the pS79F FQ-resistant strain, which exhibited a fitness advantage, was associated with a high probability of fixation (>75%). When treated with the ciprofloxacin-linezolid combination (**Figure 6B**), all other subpopulations were fully suppressed, while the two other antibiotic combinations resulted in low levels (<25%) of FQ-resistance due to the fixation of few specific subpopulations (pS79Y, pD83Y, gS81F, and gS81Y) (**Figure S7**).

The ciprofloxacin-linezolid treatment simulations informed solely by the collateral effects (Scenario 3A) showed a decrease in the emergence and fixation of pD83Y (ET 2) (**Figure 6A**), the only py strain exhibiting CS towards linezolid, compared to the simulations without any effects (Scenario 1A). All gx and py strains exhibiting CR were associated with high probability of resistance (86-97%). When such a gx strain was combined with a py strain not showing any collateral effect (*e.g.,* ET 4: gS81F and pS79F), the py-associated probability of resistance was decreased compared to when the same stain was paired with a gx showing no CR (*e.g.,* ET 3: gE85G and pS79F). Simulations of the same scenario using a combination of ciprofloxacin and penicillin showed that for all single-allele strains exhibiting CS to penicillin the probability of resistance was reduced compared to Scenario 1A. However, no evolutionary trajectories included a pair of single-allele strains where both show CS. Therefore, we are unable to elucidate if reciprocal CS could suppress all resistance development. During the free-for-all simulation, the fixation of specific FQ-resistant subpopulations differed between the treatments (**Figure 6B** **Figure S8**). However, the *de novo* emergence of single-allele subpopulations exhibiting CR was associated with high probability of resistance (>85%), while the emergence of subpopulations exhibiting CS with resistance suppression for all three antibiotic combinations.

Finally, when we compared the treatment outcome of Scenarios 1-3 to Scenario 4, with the latter taking into consideration the fitness and the collateral effects, we showed that both effects impact the probability of resistance (**Figure 6A** **Figure S7**). However, for the single-allele strains, the collateral effects seem to have a larger impact if present, as low relative growth rate combined with CR would result in high probability of resistance. The probability of the double-allele strains fixing during ciprofloxacin and erythromycin combination treatment was suppressed in the presence of CS while clearly affected by both collateral and fitness effects in the presence of CR (**Figure S7**). Overall, our findings suggest that both collateral and fitness effects can impact resistance evolution. In contrast, the simulation using the free-for-all model revealed that the fitness effect was the main driver of the resistance development during treatment with antibiotic combinations (**Figure 6B** **Figure S8**). However, due to the strong fitness advantage of pS79F observed here, which outcompete the other subpopulations, the impact of the fitness cost might be masked by the clonal suppression elicited by this subpopulation.

## Discussion

Our aim in this study was to integrate experiments and simulations to evaluate CS-informed multi-drug treatment strategies to prevent FQ-resistance in *S. pneumoni*ae. First, we characterized the collateral and fitness effects of clinically relevant FQ-resistance alleles on *S. pneumoniae* susceptibility to a wide range of antibiotics (19). We observed extensive collateral effects towards the majority of the antibiotics tested, especially antibiotics inhibiting protein synthesis, consistent with earlier results derived from laboratory-evolved strains of *Enterococcus faecalis* (7)*, Pseudomonas aeruginosa* (8, 9), *Escherichia coli* (5, 13) and *Acinetobacter baumannii* (14). CS was much more predominant than CR, in accordance with our data for this species on the collateral effects of other antibiotics (unpublished results), and the magnitude of both effects was moderate, with the exception of clindamycin and gentamicin, in line with previous findings in *E. coli* (13). Most FQ-resistant mutants also exhibited CS towards tetracycline and chloramphenicol, consistent with results in *E. faecalis* (7). FQ-resistance mutations in *gyrA* and *parC* result in three-dimensional structural changes of the DNA gyrase and the topoisomerase IV that are associated with the modification of DNA topology altering the global supercoiling of the bacterial genome (25). In *Salmonella enterica* serovar Typhimurium (26) this leads to an altered transcriptome and the subsequent global reprogramming of gene expression (27, 28), which in turn leads to collateral effects. We hypothesize that similar gene expression reprogramming, involving *i.e.*, the altered expression of stress response pathways, may also be responsible for the extensive collateral effects we observed in *S. pneumoniae*, although this remains to be elucidated in future studies.

In addition to consistent CS responses, we also observed significant heterogeneity in the CS responses of different *gyrA* and *parC* genes and alleles, even for FQ-related substitutions at single amino acids in the same gene. Most of these heterogenous effects resulted in changes to the magnitude of CS effects, but in a few cases, they also influenced the direction of the response. As yet, the cause of these differences, and of collateral effects overall, remain unclear. Because our design used transformation of gyrA and parC alleles from clinically resistant isolates of *S. pneumoniae* (19), our transformants include known QRDR mutations plus a small number of secondary mutations that are not known to be associated with FQ resistance (**Figure S9**). Although this allelic diversity reflects the natural diversity at these loci, we cannot fully exclude the influence of these non-QRDR mutations; subsequent studies using single-site mutagenesis, together with more detailed studies of transcriptional responses, will be valuable to more fully disentangle these possible influences. Overall, although our experimental results suggest that CS-informed combination treatments may be promising to limit the evolution of FQ-resistance, heterogeneous collateral effects may necessitate systematic screening of allelic identity to establish appropriate antibiotic combinations for successful selection inversion strategies.

Next, using a modeling approach, we combined our experimental findings on collateral and fitness effects, with antibiotic pharmacokinetics and pharmacodynamics in order to understand multistep resistance evolution in *S. pneumoniae* under different treatment conditions. Despite heterogeneous collateral responses for the antibiotics tested, our results simulating clinical dosing schedules for ciprofloxacin monotreatment and in combination with erythromycin, linezolid and penicillin showed a clear benefit of combination treatment over ciprofloxacin monotreatment, often successfully suppressing the emergence of FQ-resistance and clearing the infection. However, given that the clinical dosing regimens for the three antibiotics that were co-administered here with ciprofloxacin generally resulted in plasma concentration greater than the MIC of the WT (2.6 to 594 x WT MIC) and that the magnitude of the CS was generally moderate (-0.58 median fold decrease in the MIC) for these specific antibiotics, the beneficial outcome is likely driven not only by the collateral effects, but also the intrinsically increased bacterial killing associated with the addition of the second antibiotic.

*In silico* treatment combinations with average unbound steady state concentration (C_ss_) for each of the co-administered antibiotics around the MIC of the WT strain (C_ss_ 0.25-2 x MIC; far below the clinical dosing concentrations) inhibited the fixation of FQ-resistant subpopulations, and further eradicated the infection, when FQ-resistance conferred CS to the one of the co-administered antibiotics. These results indicate the potential effectiveness of CS-informed treatment combinations even with lower dosages for each of the co-administered antibiotics in eradicating infections while minimizing the risk for resistance development. This, in turn, suggests the exploitability of CS-informed treatment combinations especially for antibiotics having a narrow window between their effective doses and those doses at which they give rise to adverse toxic effects, which is in line with previous research (29).

In addition to the collateral effects, our simulations using sub-clinical dosing revealed that fitness effects had an impact on resistance development during treatment. In fact, antibiotic-resistant clonal variants encoding distinct mutations and exhibiting varying collateral and fitness effects emerge during antibiotic treatment. The clonal competition among these variants determined by their collateral and fitness effects, in turn, shapes the population dynamics and therefore the overall treatment outcome. This highlights the need to consider fitness, in addition to collateral effects, when attempting to design effective CS-informed treatments.

Translating a complex system into a simplified model requires a number of assumptions, which should be taken into consideration when interpreting our simulation results. Our pharmacodynamic model includes parameters derived from early phase data from several *in vitro* time-kill literature studies, using different *S. pneumoniae* strains (30–35). This approach allowed us to obtain typical effect parameters of *S. pneumoniae* rather than strain specific effects. We assumed that the early phase data represent the antibiotic-mediated killing of one homogenous population, thus ignoring the possibility of the observed killing rate being affected by the growth of less susceptible subpopulations. Furthermore, we assumed that there were no pharmacokinetic interactions between the antibiotics tested. Only FQ-resistance was considered in the model while resistance evolution and possible collateral effects of the second antibiotic were ignored. The model could be further expanded to include such information, thus allowing us to investigate the impact of reciprocal versus non-reciprocal collateral effects, as well as sequential treatment regimens to exploit these reciprocal effects.

In conclusion, we showed that the FQ-resistance allelic identity can have a pronounced impact on the fitness and collateral effects to different antibiotics in *S. pneumoniae* and that these can strongly influence the multistep evolution of high-level FQ-resistance under antibiotic mixing regimens. Our results highlight the importance of estimating CS for different resistance mutations to the same antibiotic; just as different mutations impart different fitness costs, so too can they drive distinct collateral responses, in turn affecting their probability of fixation during combination treatments. Despite heterogeneity in collateral effects, our results suggest that conserved CS can be exploited to develop CS-informed antibiotic treatment combinations to successfully eradicate infections while suppressing the *de novo* emergence of resistance during treatment.

## Material and Methods

### Bacterial strains, growth conditions and media

Six *S. pneumoniae* strains encoding *gyrA* and/or *parC* alleles that confer FQ-resistance were provided by the CDC Streptococcus Laboratory and used as donors for all subsequent transformations with *S. pneumoniae* D39 strain as the wild-type (WT) recipient strain. *S. pneumoniae* ATCC 49619 was used as a quality control for antimicrobial susceptibility testing. All strains used in this study and their relevant characteristics are listed in **Table S1**. Strains were routinely grown either on tryptic soy agar (BD, New Jersey, USA) supplemented with 0.5% w/v yeast extract (BD, New Jersey, USA) and 5% v/v sheep blood (Sanbio, Uden, The Netherlands) (TSYA) or in tryptic soy broth (BD, New Jersey, USA) supplemented with 0.5% w/v yeast extract (TSYB) at 37°C under 5% CO_2_ for 18-24 h, unless otherwise mentioned. The 14 antibiotics used in this study were prepared from powder stock according to manufacturers’ recommendations, stored at -20 °C or -80 °C and are listed in **Table S2**.

### Construction of fluoroquinolone-resistant mutants

Amplified fragments from genomic DNA of the FQ-resistant *S. pneumoniae* strains encoding each of the non-wild type *gyrA* or *parC* gene allele and 3 Kb of its flanking regions necessary for the integration by double crossover were used to transform the *S. pneumoniae* D39 strain. All the primer sequences used in this study are listed in **Table S3**. Transformation was performed using a saturating concentration of 1 μg/ ml of amplified DNA and 0.1 μg/ ml synthetic competence stimulating peptide 1, CSP-1 (GenScript, New Jersey, USA), as previously described (36). Putative single-allele *gyrA* (gx) and *parC* (py) mutants were selected on TSYA plates supplemented with 0.5 mg/L sparfloxacin (Santa Cruz Biotechnology, Texas, USA) or 4 mg/L ciprofloxacin (Acros Organics, New Jersey, USA), respectively. Double-allele (gxpy) mutants were generated by transforming the generated *parC* mutants with the respective *gyrA* non-WT alleles and selecting on TSYA plates supplemented with 12 mg/L ciprofloxacin (Acros Organics, New Jersey, USA). *GyrA* and *parC* allelic replacements in the transformants were confirmed by PCR and Sanger sequencing.

### Whole genome sequencing

Genomic DNA was extracted from the *S. pneumoniae* D39 WT and all FQ-resistant mutants using the DNeasy Blood & Tissue Kit (QIAGEN, Hilden, Germany) according to the manufacturer’s recommendations for Gram-positive bacteria. Quality control of purified genomic DNA was performed using Quant-iT™ dsDNA BR Assay Kit (Thermo Scientific, Breda, The Netherlands) and 1% agarose gel electrophoresis. Genomic DNA was fragmented by ultrasound on Covaris S/E210 (Covaris, Brighton, UK) according to the manufacturer’s recommendations and was subsequently used for library preparation as previously described (37). Whole genome sequencing was commercially performed using 100-bp paired-end libraries on a BGISEQ-500 platform at BGI Tech Solutions (Hong Kong, China) or on a NextSeq 2000 platform at MiGS (Pittsburgh, USA). High-quality filtered reads were mapped to the reference genome of *S. pneumoniae* D39V available in GenBank (accession number: NZ_CP027540.1) in order to identify and annotate genetic differences found between our FQ-resistant mutants and their parental D39 WT strain using the open-source computational pipeline *breseq* with the default parameter settings (38).

### Antimicrobial susceptibility testing

Minimum inhibitory concentrations (MICs) of all isogenic strains for 12 clinically relevant antibiotics and ciprofloxacin (**Table S2**) were determined in triplicate by broth microdilution according to European Committee on Antimicrobial Susceptibility Testing (EUCAST; http://www.eucast.org) and ISO 20776-1:2006 guidelines with two minor modifications. A 1.5-fold testing scale was used to include the standard two-fold antibiotic concentrations and their median values and the cation-adjusted Mueller-Hinton broth (CAMHB - BD, New Jersey, USA) was supplemented with 100 U of catalase (Worthington Biochemical Corporation, New Jersey, USA) instead of 5% lysed horse blood. The lowest antibiotic concentration where no turbidity was observed was scored as the MIC.

### Static time-kill assays

Static time-kill assays were performed in triplicate using starting inoculum of the parental D39 WT strain equating to ∼10^6^ CFU/mL in 15 mL of CAMHB supplemented with 5% lysed blood according to EUCAST and ISO 20776-1:2006 guidelines. Antibiotic-free cultures and cultures with either the addition of antibiotics as monotherapy or in combinations at final concentrations equal to the average unbound steady state concentration (C_ss_) of each antibiotic were incubated at 37°C under 5% CO_2_ continuous agitation (100 rpm). At set time points of 0, 3, and 6 h post inoculation, 100 μL samples were collected, serially diluted and plated on TSYA plates. After 48 h of incubation at 37°C and 5% CO_2_, colonies were counted for viable cell titer determination.

### Collateral effect determination

Collateral effects were determined as the decrease (CS) or increase (collateral resistance, CR) in the MIC of each mutant relative to the isogenic WT D39 strain, and their magnitude as the log2-scaled fold change in the MICs between each mutant and the WT D39 strain (Equation 1), as previously described (7).

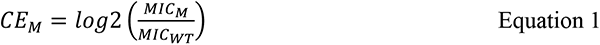

Where CE_M_ represents the collateral effect of a mutant (gx, py, or gxpy), MIC_M_ its corresponding MIC and MIC_WT_ the MIC of the D39 WT strain.

The conservation of the collateral effects was assessed based on the CS_50_ and CR_50_ thresholds, defined respectively as CS and CR effects occurring in more than 50% of the mutants tested (13).

### Growth rate measurements

The *in vitro* fitness of the WT and each mutant was determined in triplicate using the Malthusian growth model (39). Bacterial inocula of ∼10^8^ CFU/ ml were diluted 100-fold into 3 ml of fresh pre-warmed TSYB medium. 200 µl of each diluted culture was loaded in wells of a 100-well honeycomb plate and incubated at 37 °C. Subsequently, the optical density at 600 nm was measured hourly using a Bioscreen C Reader (Thermo Scientific, Breda, The Netherlands), with 5 s of shaking before reads. Growth rates were calculated based on the slope of the line that fitted points displaying log-linear growth. Relative growth rates were calculated by dividing the generation time of each D39 mutant by that of the mean growth rate of the WT D39 strain.

### Statistical analysis

Statistical analyses were performed using R (version 3.6.3). We performed hierarchical clustering to identify relationships between strains, and between antibiotics, relating to collateral effects using the complete linkage method within the stats R package. We evaluated between and within group differences in mean relative growth rate with a one-way ANOVA and a post-hoc Tukey pairwise comparison with multiple correction when significant differences were identified (α = 0.05).

### Mathematical modeling

We performed mathematical modeling of bacterial growth dynamics to evaluate the effect of different FQ mono- and combination-treatments (antibiotic mixing) for eight different trajectories leading to high-level FQ-resistance on treatment outcomes and the probability of resistance. The model included clinical pharmacokinetics of antibiotics used in the treatment schedule, bacterial growth rates, antibiotic-mediated killing, collateral effects, and stepwise FQ-resistance development (**Figure 3B**). The treatment simulations were conducted for ciprofloxacin monotreatment and in combination with erythromycin, linezolid, and penicillin, which were chosen because of the availability of literature derived experimental data (30–35) needed for the estimation of pharmacodynamic parameters. We incorporated the experimentally measured relative fitness and collateral effects observed for the strains designed in this study.

The bacterial model was comprised of a four-state stochastic hybrid ordinary differential equation (ODE) model, where each state represents a bacterial subpopulation. Included *S. pneumoniae* subpopulations were the FQ-sensitive D39 wild-type (WT; *gyrA*_wt_::*parC*_wt_), the resistant *gyrA* mutant (gx; *gyrA*_x_::*parC*_wt_), the resistant *parC* mutant (py; *gyrA*_wt_::*parC_y_*), and the resistant *gyrA* and *parC* double mutant (gxpy; *gyrA*_x_::*parC_y_*). Evolution of *de novo* resistance occurred in a stepwise manner with the emergence of either of the low-level FQ-resistant single-allele mutants (gx or py), followed by the emergence of their corresponding high-level FQ-resistant double-allele mutant (gxpy). The mutation events were modelled by a stochastic process calculating the specific number of bacteria mutated for each time interval *τ*, which was converted to a transition- and interval-specific mutation rate (*k_z,M,τ_*) using a τ of one hour. This mutation process was based on a binomial distribution *B* with a mutation probability equal to the *gyrA* and *parC* mutation frequency per hour (μ) (40), as shown in Equation 2, and was identically implemented for each state transition.

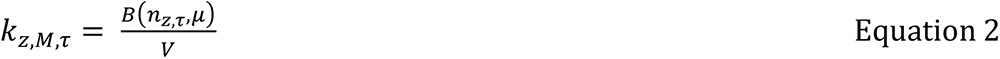

where *n_z,τ_* is the integer number of bacteria of subpopulation *z* at start of τ and *V* is the volume of the infection site.

Each bacterial state (*z*) included a logistic growth model for which the state-specific maximal growth rate (*K_G,z_*) was a composite parameter of the literature derived growth rate of D39 (41) and our experimental growth rate ratio between mutant and WT. The systems growth capacity limitation parameter (*Bmax*) and the bacterial concentration at the start of the infection were obtained from an *in vivo* model of bacteraemia (24). The volume available for infection represented a human blood volume (42). All system specific parameters are stated in **Table S4**.

Antibiotic-mediated killing was implemented for each state *z* (*E_AB,_*) according to Equation 3, which includes the subpopulation-specific antibiotic effect of AB_1_ (*E_AB_1_,z_*), AB_2_ (*E_AB_2_,z_*), and a term describing the PD interaction between these drugs 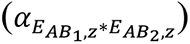, where significant interactions were identified based on our performed static time-kill assays.

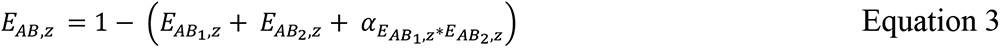

The antibiotic-specific effects were described according to a previously described model (43) where the effect of the *i*^th^ antibiotic on bacterial state *z* (*E_AB_i_,z_*) related to the unbound concentration (*C_AB,*i*_*) according to Equation 4.

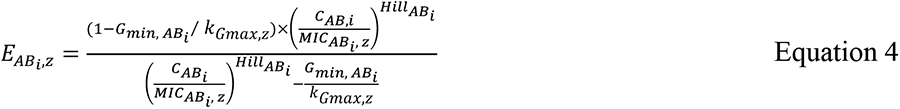

Where *G_min,AB_*i*__* represents the maximal killing rate, *Hill_AB_i__* the shape factor of the concentration-effect relationship, *k_Gmax,z_* the state specific maximal growth rate, and *MIC_AB_i_,z_* the state specific MIC.

Pharmacodynamic model parameter values were obtained by fitting the model to digitized early phase *S. pneumoniae* experimental *in vitro* time-kill data using the nlmixr package (version 1.1.1-7) within the statistical software R (version 3.6.3). Growth in the absence of antibiotic and antibiotic-mediated bacterial killing were modelled sequentially. Initially, bacterial maximal growth rates were obtained by fitting an exponential growth model for all the antibiotic-free growth data. A log-normal random effect was included on maximal growth rate to account for between experiment variability. The estimated individual maximal growth rates were included as a covariate in the subsequent fitting of the antibiotic-mediated killing. Separate models were fitted for each antibiotic to obtain the drug specific parameters *E_max,AB_i__*. and *hill_AB_i__*. Antibiotic-mediated killing was incorporated separately for each antibiotic on each individual bacterial state with the corresponding experimentally determined MICs (*MIC_AB_i_,z_*).

Data generated through the interaction study were fitted with non-linear models including exponential bacterial growth and antibiotic-mediated killing implemented according to Equation 5.

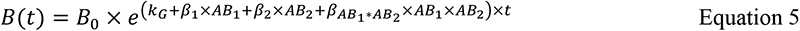

Where *B* represents the bacterial density, *B_0_* the inoculum, *k_G_* the bacterial maximal growth rate, *β_1_* the effect size of antibiotic 1 (AB_1_), *β_2_* the effect size of antibiotic 2 (AB_2_), and *β_AB_1_*AB_2__* the effect size of the interaction between AB_1_ and AB_2_. To facilitate the translation of the integration to the PK-PD model framework, a relative interaction term (*θ_AB_1_*AB_2__*) was derived according to Equation 6.

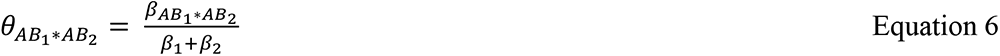

The PD interaction was subsequently integrated to the framework according to Equation 7.

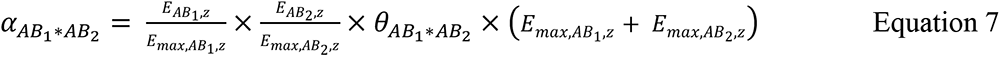

Where *E_max, AB_i_,z_* is given by Equation 8.

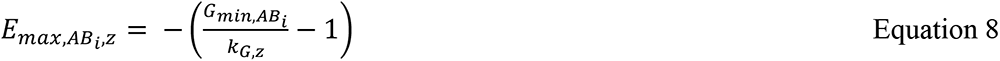

The subpopulation-specific rates of change for bacterial density are shown in the Equation 9-12. These rates are dependent on the bacterial density of the specific subpopulation, the subpopulation-specific net growth (*k_net,z_*), the antibiotic effect (*E_AB,z_*), and the mutational transition(s) (*k_z_*,*M,τ*), if present.

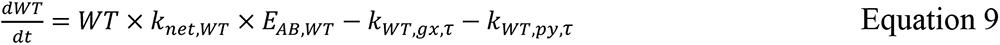

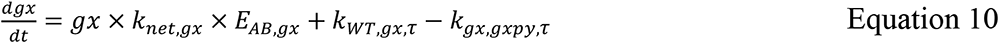

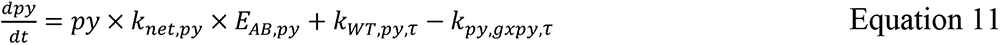

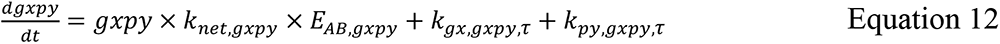

Where *k_net,z_* is given by Equation 13.

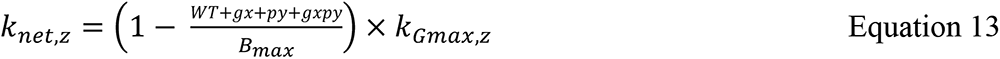

The pharmacodynamic model was linked to a previously published pharmacokinetic model of ciprofloxacin (44) and a pharmacokinetic model of the respective antibiotic selected for combination treatment (**Table S5**). The pharmacokinetic models were used to simulate pharmacokinetic profiles of an individual with a body weight of 70 kg and a creatinine clearance of 75 mL/min/1.73 m^2^. The estimate of the 95^th^ percentile of parameters related to clearance and volume were used if the inter-individual variability was reported. The fraction unbound (f_u_) was used to obtain the unbound concentration of each antibiotic.

The modeling framework was used to simulate two different sets of treatment scenarios of *S. pneumoniae* bacteraemia: 1) using ciprofloxacin as mono-treatment (500 mg twice daily (b.i.d.)) or in combination with erythromycin (600 mg b.i.d.), linezolid (600 mg b.i.d.), or penicillin (3 g b.i.d.); and 2) using a series of non-clinical treatment scenarios where the average unbound steady state concentration (C_ss_) of each antibiotic was related to the MIC of the WT (C_ss_ 0.25-2 x MIC). A third set of simulations was performed where we investigated and disentangled the impact of fitness and collateral effect on resistance development by removing one or both of the experimentally derived effects. For these simulations, we focused on the concentrations associated with large between-trajectory variability in the previous set of simulations. The selection of the anti-pneumococcal antibiotics included in these treatment scenarios was based on the availability of relevant time-kill data and ciprofloxacin was used as representative agent of the FQ class. Each of the treatment scenarios was simulated 500 times. The treatment outcomes were evaluated by assessing the probability of failure in eradicating the WT strain and/or the probability of resistance establishment. The failure in WT eradication and resistance establishment were defined respectively as a bacterial density of WT and FQ-resistant mutant strains exceeding 10^4^ CFU/ml at the end of treatment (two weeks), with the particular density corresponding to both the initial inoculum and the density of an established infection (24). The framework was used to simulate treatments of eight defined trajectories, each depicting the *de novo* multistep emergence of a particular high-level FQ-resistant double-allele mutant (gxpy), with either of its corresponding low-level FQ-resistant single-allele mutants (gx or py) as an intermediate step (**Figure 2**).

The modeling framework was adapted to allow for a free-for-all simulation, where all evolutionary trajectories were included simultaneously. Thus, the adapted model constituted of four gx subpopulations, four py subpopulations, and eight gxpy subpopulations. Appropriate state transitions were included on gx and py subsequently leading to the eight gxpy studied. This adapted model was used to further investigate the role of the collateral and fitness effects in the context of greater clonal competition.

### Software and model code

All model simulations were conducted in R (version 3.6.3), using the ODE solver package RxODE (version 1.0.0). The modeling framework, the adapted free-for-all version and the associated code are available at Github (https://github.com/vanhasseltlab/PKPD-CSpneumo) (50).

## Acknowledgements

We are thankful to Dr. Lesley McGee and the CDC Streptococcus Laboratory for providing us with the clinical strains carrying the relevant *gyrA* and *parC* alleles conferring fluoroquinolone-resistance. We are, also, thankful to Dr. Bastienne Vriesendorp for her help on the SNP calling. We acknowledge the helpful comments of Dr. Pål J. Johnsen on an earlier version of this manuscript and the valuable input of Dr. Shraddha Shitut on the design of Figure 1A of this manuscript. AL and DER were supported through the JPI-EC-AMR (Project 547001002). JGCvH was supported by ZonMW Off Road (Project 451001033).

## Notes

### Competing Interest Statement

The authors have declared no competing interest.

