## Supplementary Material for "Allele-specific collateral and fitness effects determine the dynamics of fluoroquinolone-resistance evolution"

### Supplementary Tables

**Table S1.** *Streptococcus pneumoniae* strains included in this study and their relevant characteristics.

| **Strains** | **Relevant characteristic(s)** | **Source** |
|---|---|---|
| D39 | Encapsulated, Serotype 2, wild-type *gyrA* and *parC* | Laboratory stock |
| ATCC 49619 | Encapsulated, Serotype 19F, wild-type *gyrA* and *parC* | Laboratory stock |
| SP12 | FQ-resistant ($gyrA_{S81Y}$), donors for transformations | CDC |
| SP35 | FQ-resistant ($gyrA_{S81F}$::$parC_{D83Y}$), donors for transformations | CDC |
| SP47 | FQ-resistant ($gyrA_{S81F}$::$parC_{D83N}$), donors for transformations | CDC |
| SP60 | FQ-resistant ($gyrA_{S81F}$::$parC_{S79F}$), donors for transformations | CDC |
| SP62 | FQ-resistant ($gyrA_{E85G}$::$parC_{S79F}$), donors for transformations | CDC |
| SP95 | FQ-resistant ($gyrA_{E85K}$::$parC_{S79Y}$), donors for transformations | CDC |
| MgS81F | FQ-resistant ($gyrA_{S81F}$::$parC_{WT}$), isogenic D39 transformant | This study |
| MgS81Y | FQ-resistant ($gyrA_{S81Y}$::$parC_{WT}$), isogenic D39 transformant | This study |
| MgE85G | FQ-resistant ($gyrA_{E85G}$::$parC_{WT}$), isogenic D39 transformant | This study |

| | | |
|---|---|---|
| MgE85K | FQ-resistant (*gyrA*$_{E85K}$::*parC*$_{WT}$), isogenic D39 transformant | This study |
| MpS79F | FQ-resistant (*gyrA*$_{WT}$::*parC*$_{S79F}$), isogenic D39 transformant | This study |
| MpS79Y | FQ-resistant (*gyrA*$_{WT}$::*parC*$_{S79Y}$), isogenic D39 transformant | This study |
| MpD83N | FQ-resistant (*gyrA*$_{WT}$::*parC*$_{D83N}$), isogenic D39 transformant | This study |
| MpD83Y | FQ-resistant (*gyrA*$_{WT}$::*parC*$_{D83Y}$), isogenic D39 transformant | This study |
| MgS81F::pS79F | FQ-resistant (*gyrA*$_{S81F}$::*parC*$_{S79F}$), isogenic D39 transformant | This study |
| MgS81F::pS79Y | FQ-resistant (*gyrA*$_{S81F}$::*parC*$_{S79Y}$), isogenic D39 transformant | This study |
| MgS81F::pD83N | FQ-resistant (*gyrA*$_{S81F}$::*parC*$_{D83N}$), isogenic D39 transformant | This study |
| MgS81F::pD83Y | FQ-resistant (*gyrA*$_{S81F}$::*parC*$_{D83Y}$), isogenic D39 transformant | This study |
| MgS81Y::pS79F | FQ-resistant (*gyrA*$_{S81Y}$::*parC*$_{S79F}$), isogenic D39 transformant | This study |
| MgS81Y::pS79Y | FQ-resistant (*gyrA*$_{S81Y}$::*parC*$_{S79Y}$), isogenic D39 transformant | This study |
| MgE85G::pS79F | FQ-resistant (*gyrA*$_{E85G}$::*parC*$_{S79F}$), isogenic D39 transformant | This study |
| MgE85K::pS79Y | FQ-resistant (*gyrA*$_{E85K}$::*parC*$_{S79Y}$), isogenic D39 transformant | This study |

**Table S2.** List of antibiotics used in this study and their targets.

| **Antibiotic Name (Abbreviation)** | **Antibiotic class** | **Antibiotic target(s)** | **Supplier** |
|---|---|---|---|
| Chloramphenicol (CHL) | Amphenicol | Protein synthesis (50S) | Sigma-Aldrich |
| Ciprofloxacin (CIP) | Quinolone | DNA replication (GyrA + ParC) | Acros Organics |
| Clindamycin (CLI) | Lincosamide | Protein synthesis (50S) | Cayman chemical |
| Daptomycin (DAP) | Lipopeptide | Cell membrane | Acros Organics |
| Erythromycin (ERY) | Macrolide | Protein synthesis (50S) | Sigma-Aldrich |
| Fusidic Acid (FUS) | Fusidane | Protein synthesis (EF-G) | Cayman chemical |
| Gentamicin (GEN) | Aminoglycosides | Protein synthesis (30S) | Sigma-Aldrich |
| Linezolid (LNZ) | Oxazolidinone | Protein synthesis (50S) | Cayman chemical |
| Penicillin (PEN) | β-lactam | Cell wall synthesis (PBPs) | Sigma-Aldrich |
| Rifampicin (RIF) | Rifamycin | RNA synthesis (rpoB) | GERBU Biotechnik |
| Sparfloxacin (SPR) | Quinolone | DNA replication (GyrA + ParC) | Cayman chemical |

| | | | |
|---|---|---|---|
| Trimethoprim/sulfamethoxazole (SXT) | Antifolate | Folate synthesis (FolA + FolP) | SERVA/ AG Scientific |
| Tetracycline (TET) | Tetracycline | Protein synthesis (30S) | Sigma-Aldrich |
| Vancomycin (VAN) | Glycopeptide | Cell wall synthesis | Carl Roth |

**Table S3.** Oligonucleotides used in this study.

| Oligonucleotide | Sequence (5′-3′) | Purpose | Reference |
|---|---|---|---|
| GyrA_3kb_F | AATTATCAACATCGACAAAGG | Amplification of *gyrA* gene and ~ 3Kb flanking regions for allelic replacement. | This study |
| GyrA_3kb_R | AACACTTGAGAATGAAATTCG | Amplification of *gyrA* gene and ~ 3Kb flanking regions for allelic replacement. | This study |
| ParC_3kb_F | ACGAATGATAATCAAACTAGC | Amplification of *parC* gene and ~ 3Kb flanking regions for allelic replacement. | This study |
| ParC_3kb_R | CTAAATTCTGGAATCATTTGC | Amplification of *parC* gene and ~ 3Kb flanking regions for allelic replacement. | This study |
| gyrA-F-seq | GATACCTGGCTAACTTGATA | Sequencing of *gyrA* gene for confirmation of allelic replacement. | This study |
| gyrA-R-seq | GATTTTCCCTAGACAACTTC | Sequencing of *gyrA* gene for confirmation of allelic replacement. | This study |
| parC-F-seq | CCTACTCTACATTCTTTGAAA | Sequencing of *parC* gene for confirmation | This study |

| | | | |
|---|---|---|---|
| | | of allelic replacement. | |
| parC-R-seq | TTACTGTCATATTCCACTCC | Sequencing of *parC* gene for confirmation of allelic replacement. | This study |

**Table S4.** System specific model parameters.

| Parameter | Value | Note | Reference |
|---|---|---|---|
| Growth rate ($k_{G,z}$) | 0.77 $h^{-1}$ *ratio | Composite parameter of literature D39 natural growth and experimental ratio between WT and mutant | (40) |
| Mutation rate ($\mu$) | 1.4 x $10^{-8}$ | Literature value for D39 *gyrA* and *parC* mutations. The resistance frequency was calculated as the number of resistant colonies per inoculum (~$10^8$ cells). | (39) |
| Maximal capacity ($B_{max}$) | $10^8$ cfu/ml | Maximal capacity in cfu/ml derived from blood sampling of *in vivo* experiment day 4 | (24) |
| Inoculum | $10^4$ cfu/ml | Derived from blood sampling of *in vivo* experiment day one | (24) |
| Infection volume | 5 L | Volume of blood to represent bacteraemia | (41) |

**Table S5.** Pharmacokinetic model parameters of the antibiotics selected for combination treatment.

| Description | Parameter | Ciprofloxacin | Erythromycin | Linezolid | Penicillin |
|---|---|---|---|---|---|
| Central volume | Vc | 23.6 L | 34.9 L | 67.2 L | 18.1 L |
| Clearance | CL | 82.9 L/h | 22.8 L/h | 5.8 L/h | 34.3 L/h |
| Peripheral volume | Vp | NA | 20.7 L | NA | NA |
| Inter-compartmental clearance | Q | NA | 16.9 L/h | NA | NA |
| Unbound fraction | fu | 0.79 (44) | 0.16 | 0.815 (45) | 0.40 |
| Reference | | (43) | (46) | (47) | (48) |

All models were either one- or two-compartmental models with first order elimination.

NA=Not applicable.

**Supplementary Figures**

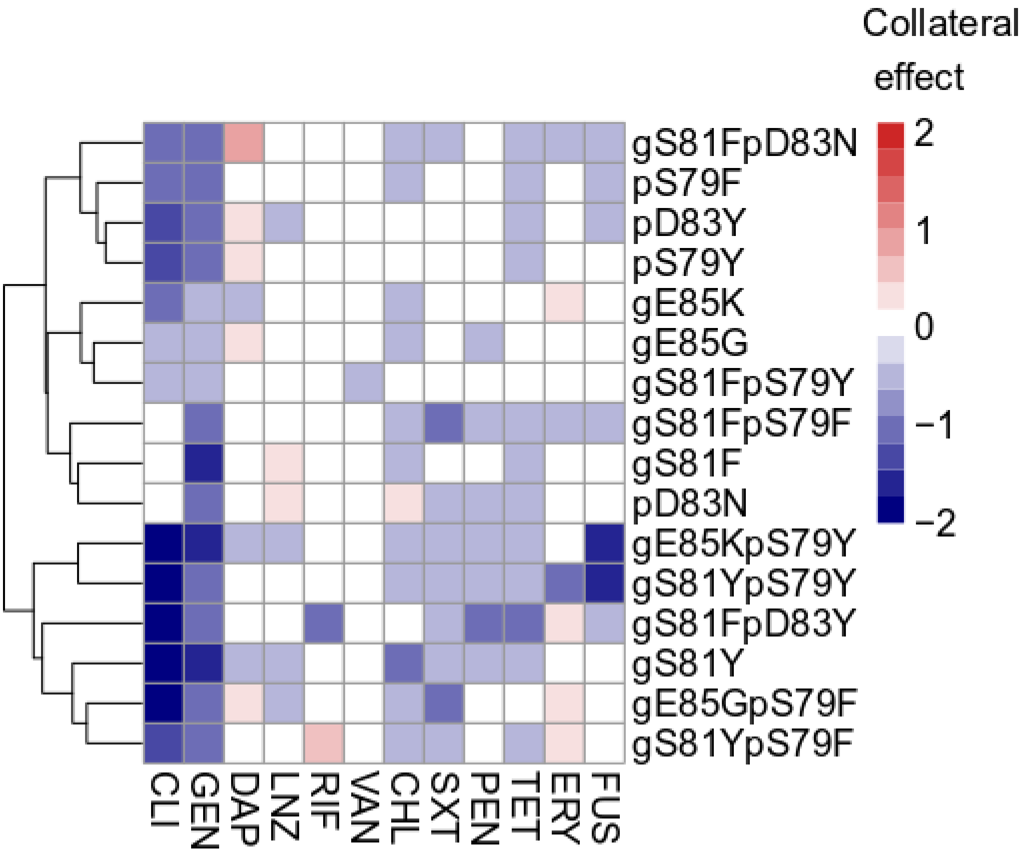


**Figure S1.** Hierarchical clustering based on collateral effects between strains and antibiotics. Collateral effect for FQ-resistant strains (y axis) against different antibiotics (x-axis). Color intensity corresponds to the collateral effect magnitude quantified by the mean log2 relative change of MIC compared to the WT. Collateral resistance (CR) and collateral sensitivity (CS) are respectively depicted with red and blue.

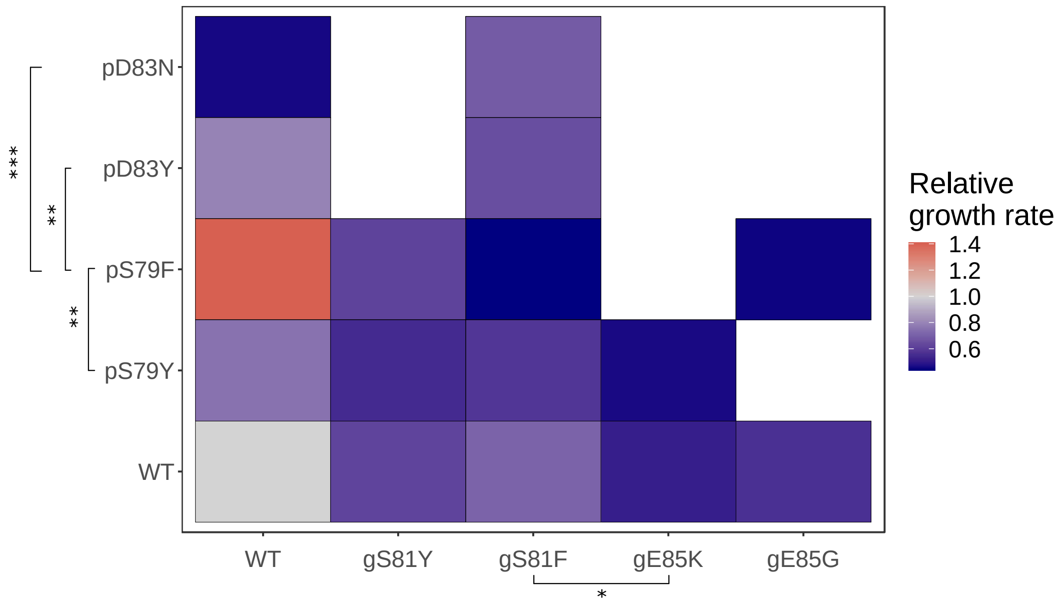


**Figure S2.** Fitness of FQ-resistant *S. pneumoniae*. **A:** Mean relative growth rate of three replicates of mutants harboring specific *gyrA* (gx) and/or *parC* (py) mutations. Significant differences are indicated by * (p<0.05), ** (p<0.01), *** (p<0.001) and **** (p <0.0001).

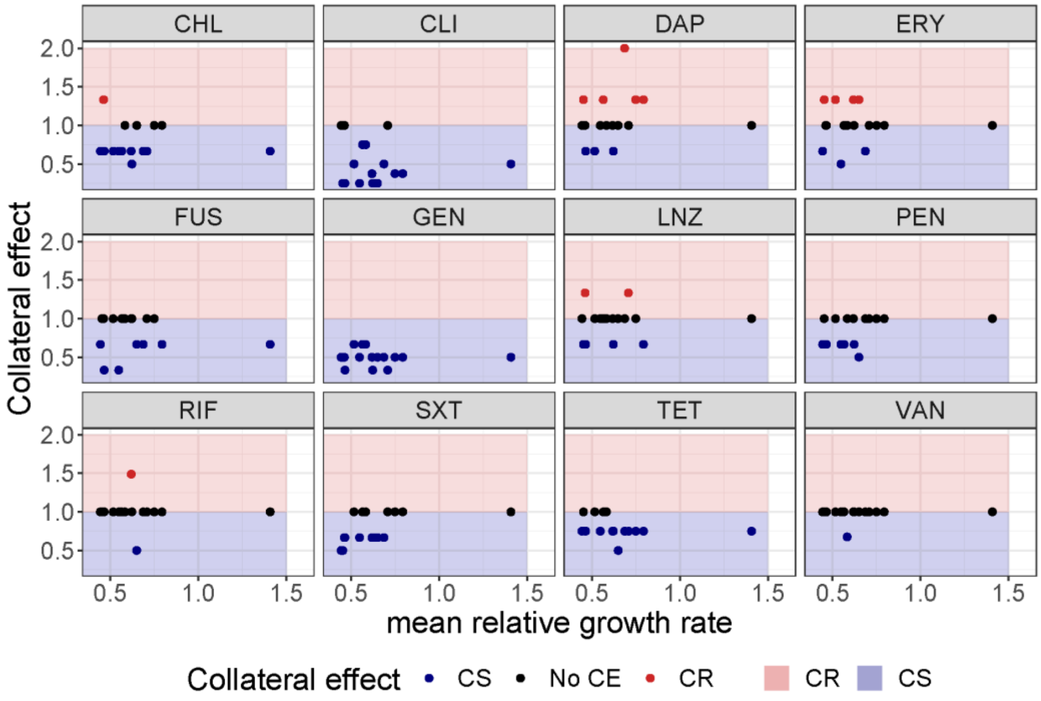


**Figure S3.** Correlation between mean relative growth rate (KG) and collateral effect (CE). Mean relative growth rate of mutants encoding specific FQ-resistant *gyrA* (gx) and/or *parC* (py) alleles compared to WT versus CE, where red indicate collateral resistance (CR), black no CE, and blue collateral sensitivity (CS).

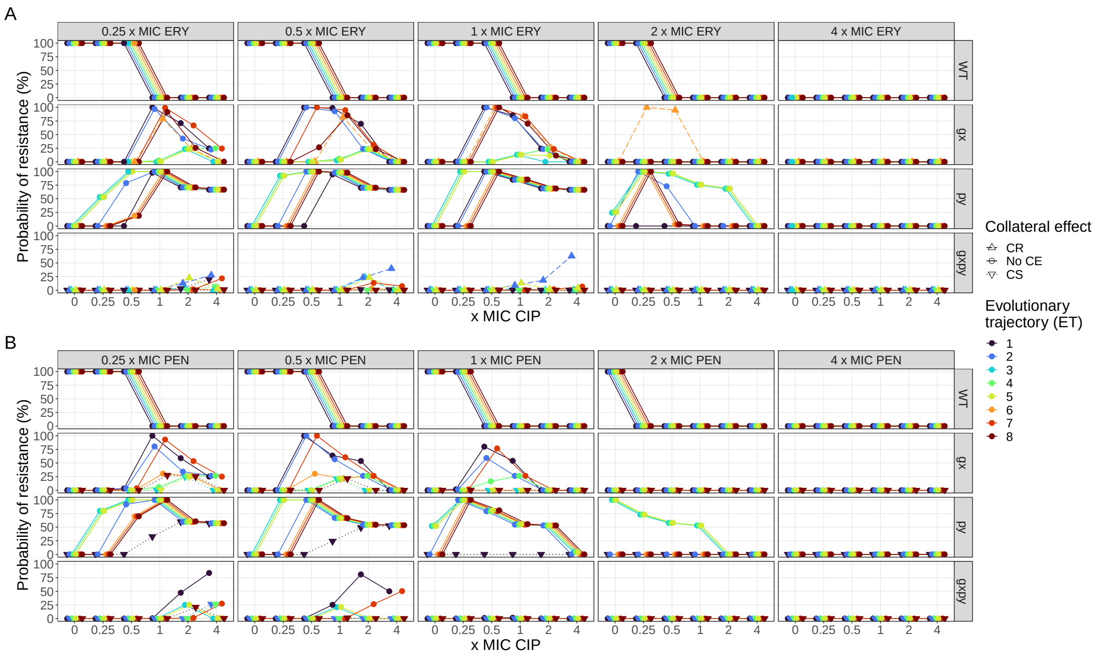


**Figure S4.** Treatment outcome of ciprofloxacin (CIP) in combination with erythromycin (ERY) or penicillin (PEN). The treatments were evaluated on eight different trajectories leading to high-level FQ-resistance (see Figure 4A). Simulation outcomes of CIP in combination with ERY (**A**) or PEN (**B**) with average unbound steady state concentrations ($C_{ss}$) around the MIC of the WT population ($C_{ss}$ 0.25 - 4 x MIC). Each point represents a bacterial subpopulation from a specific trajectory (ET, indicated by color). Collateral effects (CE) are indicated by shape and line-type, where solid lines with round points represents no CE, dashed lines with triangles pointing up collateral resistance (CR), and dotted lines with triangles pointing down collateral sensitivity (CS).

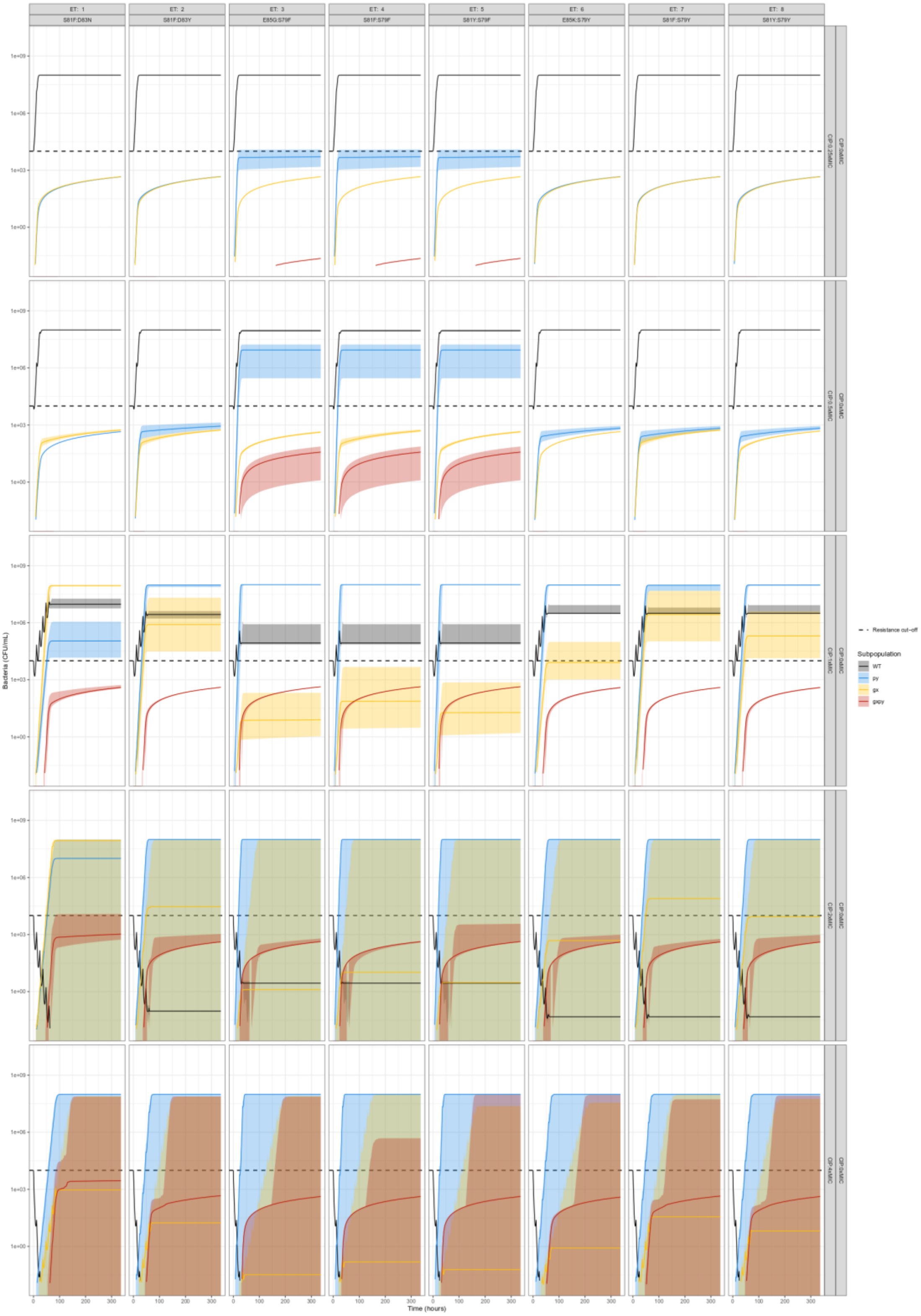
ET: 1
S81F:D83N
ET: 2
S81F:D83Y
ET: 3
E85G:S79F
ET: 4
S81F:S79F
ET: 5
S81Y:S79F
ET: 6
E85K:S79Y
ET: 7
S81F:S79Y
ET: 8
S81Y:S79Y
CIP:0.25xMIC
CIP:0.5xMIC
CIP:1xMIC
CIP:2xMIC
CIP:4xMIC
CIP:0xMIC
1e+09
1e+06
1e+03
1e+00
Bacteria (CFU/mL)
0
100
200
300
Time (hours)
Resistance cut-off
Subpopulation
WT
py
gx
gxpy

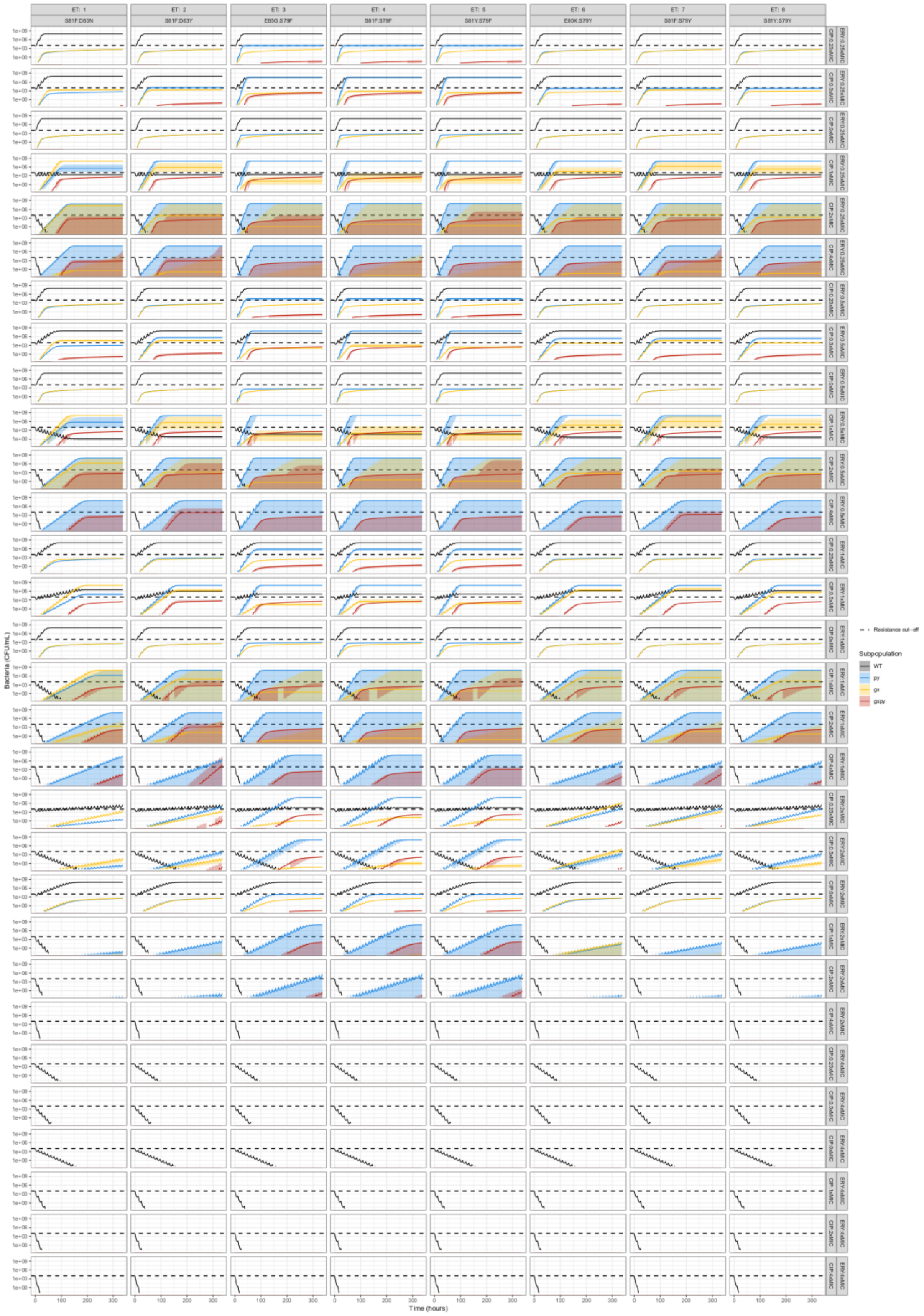

ET: 1
S81F:D83N
ET: 2
S81F:D83Y
ET: 3
E85G:S79F
ET: 4
S81F:S79F
ET: 5
S81Y:S79F
ET: 6
E85K:S79Y
ET: 7
S81F:S79Y
ET: 8
S81Y:S79Y
ERY:0.25xMIC
CIP:0.25xMIC
Bacteria (CFU/mL)
Time (hours)
Resistance cut−off
Subpopulation
WT
py
gx
gxpy

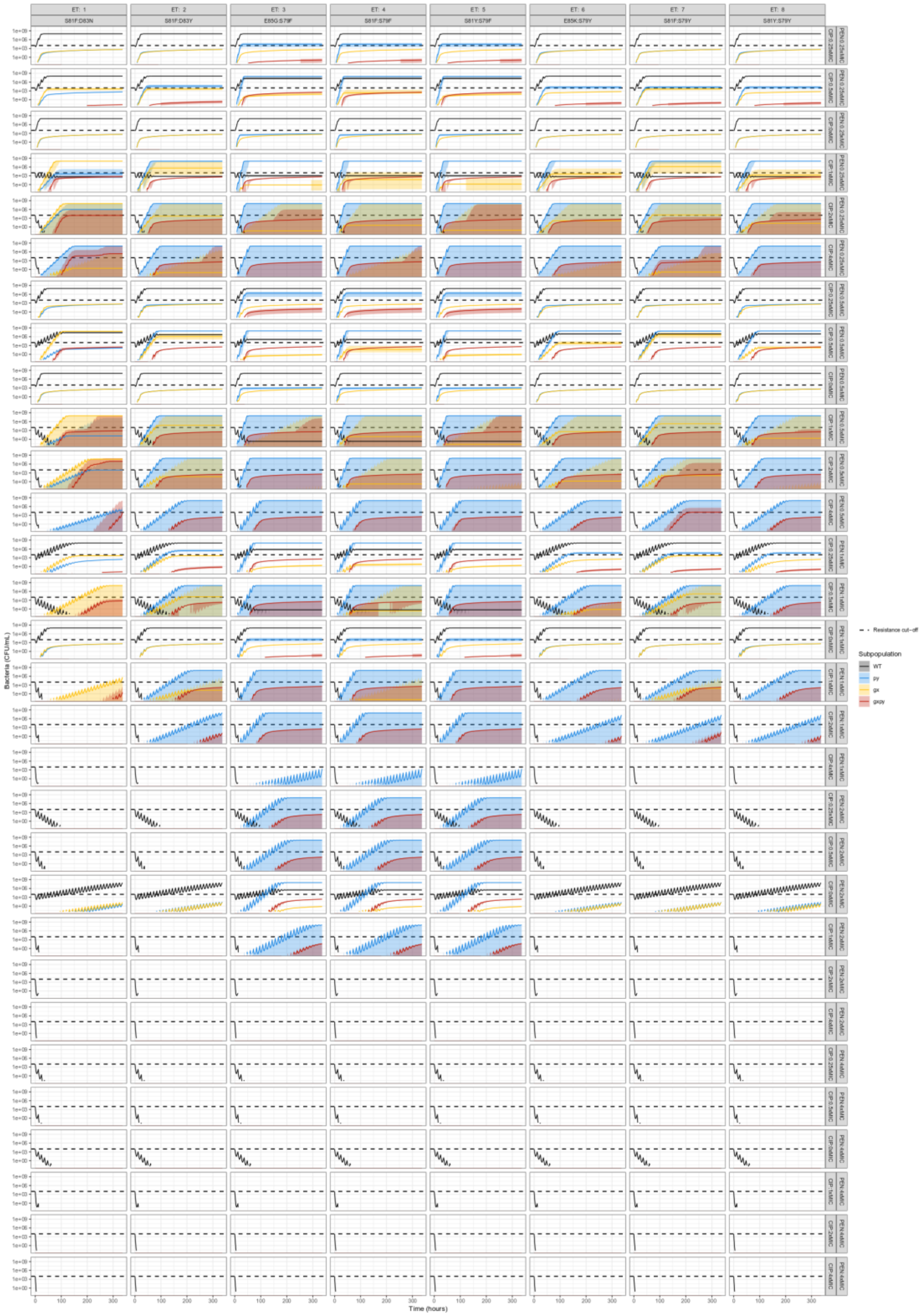
ET: 1
S81F:D83N
ET: 2
S81F:D83Y
ET: 3
E85G:S79F
ET: 4
S81F:S79F
ET: 5
S81Y:S79F
ET: 6
E85K:S79Y
ET: 7
S81F:S79Y
ET: 8
S81Y:S79Y
Bacteria (CFU/mL)
1e+09
1e+06
1e+03
1e+00
Time (hours)
0
100
200
300
Resistance cut-off
Subpopulation
WT
py
gx
gxpy

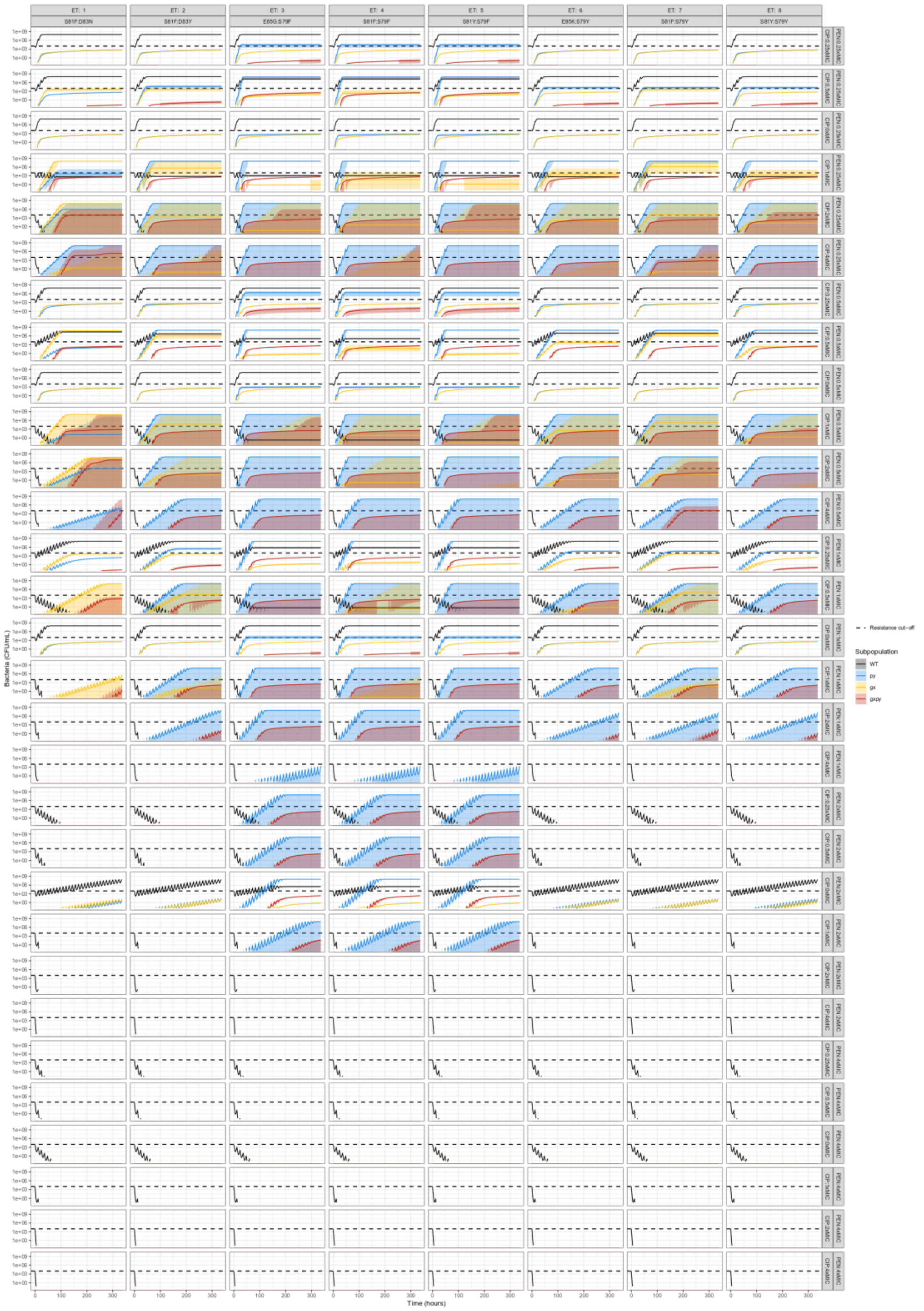


**Figure S5.** Overall bacterial dynamics of simulated ciprofloxacin mono- or combination with erythromycin (ERY), linezolid (LNZ), or penicillin (PEN). Each simulation includes WT (green), a *gyrA* mutant gx (yellow), a *parC* mutant py (light blue), and the double-allele mutant gxpy (red). Simulation outcomes treated with CIP in combination with ERY, LNZ, or PEN with average unbound steady state concentrations ($C_{ss}$) around the MIC of the WT population ($C_{ss}$ 0.25 - 4 x MIC). The treatments were evaluated on eight different trajectories leading to high-level FQ-resistance (see Figure 5A). Solid line represents the median bacterial growth dynamics of 1000 simulations where bacterial subpopulations are indicated by the respective color. The shaded areas represent the 5$^{th}$ and 95$^{th}$ simulated percentiles. The black dashed line represents the resistance cut-off, which is equal to the WT bacterial density at start of treatment.

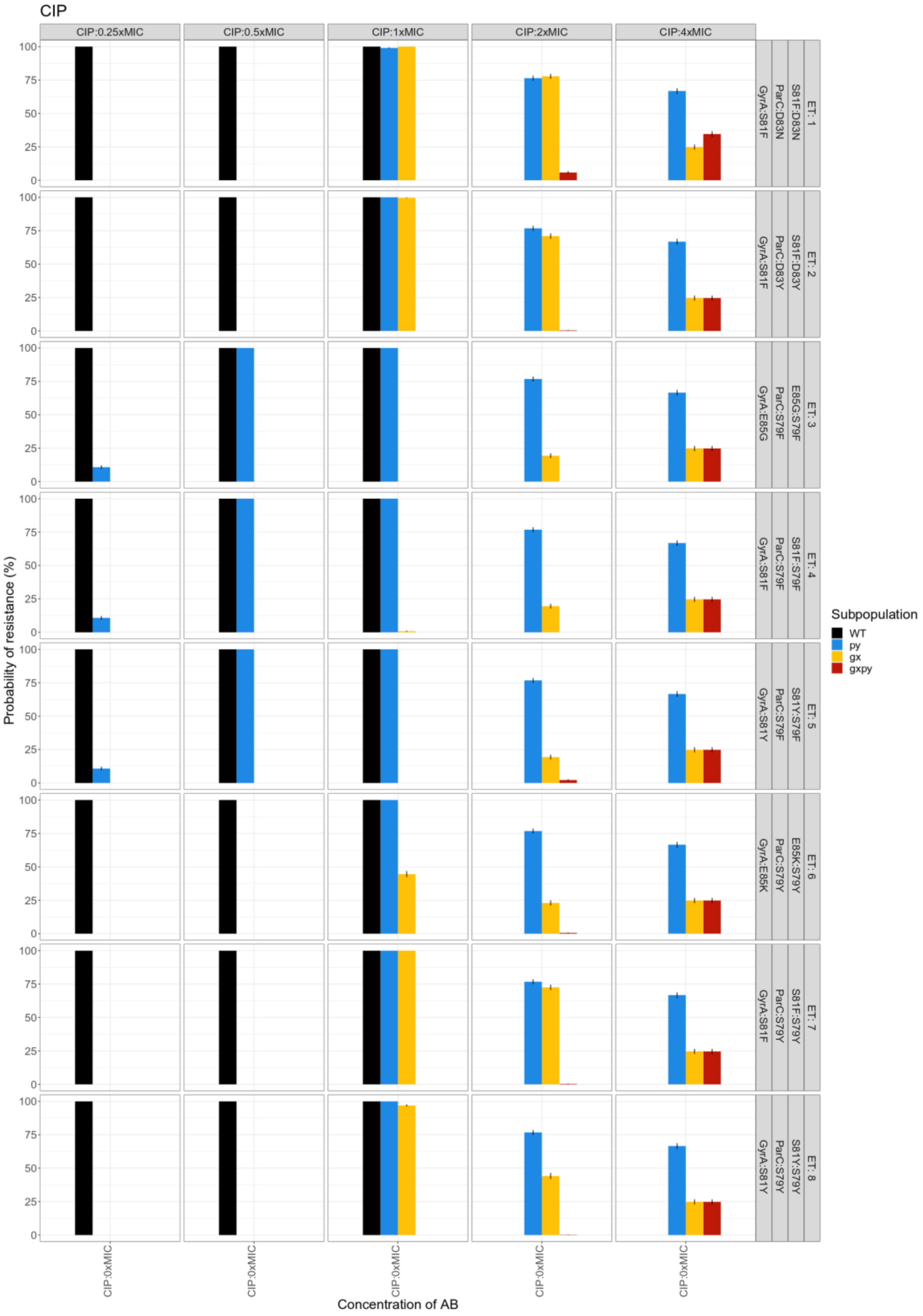
CIP
CIP:0.25xMIC
CIP:0.5xMIC
CIP:1xMIC
CIP:2xMIC
CIP:4xMIC
ET: 1 S81F:D83N ParC:D83N GyrA:S81F
ET: 2 S81F:D83Y ParC:D83Y GyrA:S81F
ET: 3 E85G:S79F ParC:S79F GyrA:E85G
ET: 4 S81F:S79F ParC:S79F GyrA:S81F
ET: 5 S81Y:S79F ParC:S79F GyrA:S81Y
ET: 6 E85K:S79Y ParC:S79Y GyrA:E85K
ET: 7 S81F:S79Y ParC:S79Y GyrA:S81F
ET: 8 S81Y:S79Y ParC:S79Y GyrA:S81Y
Probability of resistance (%)
Concentration of AB
CIP:0xMIC
Subpopulation
WT
py
gx
gxpy

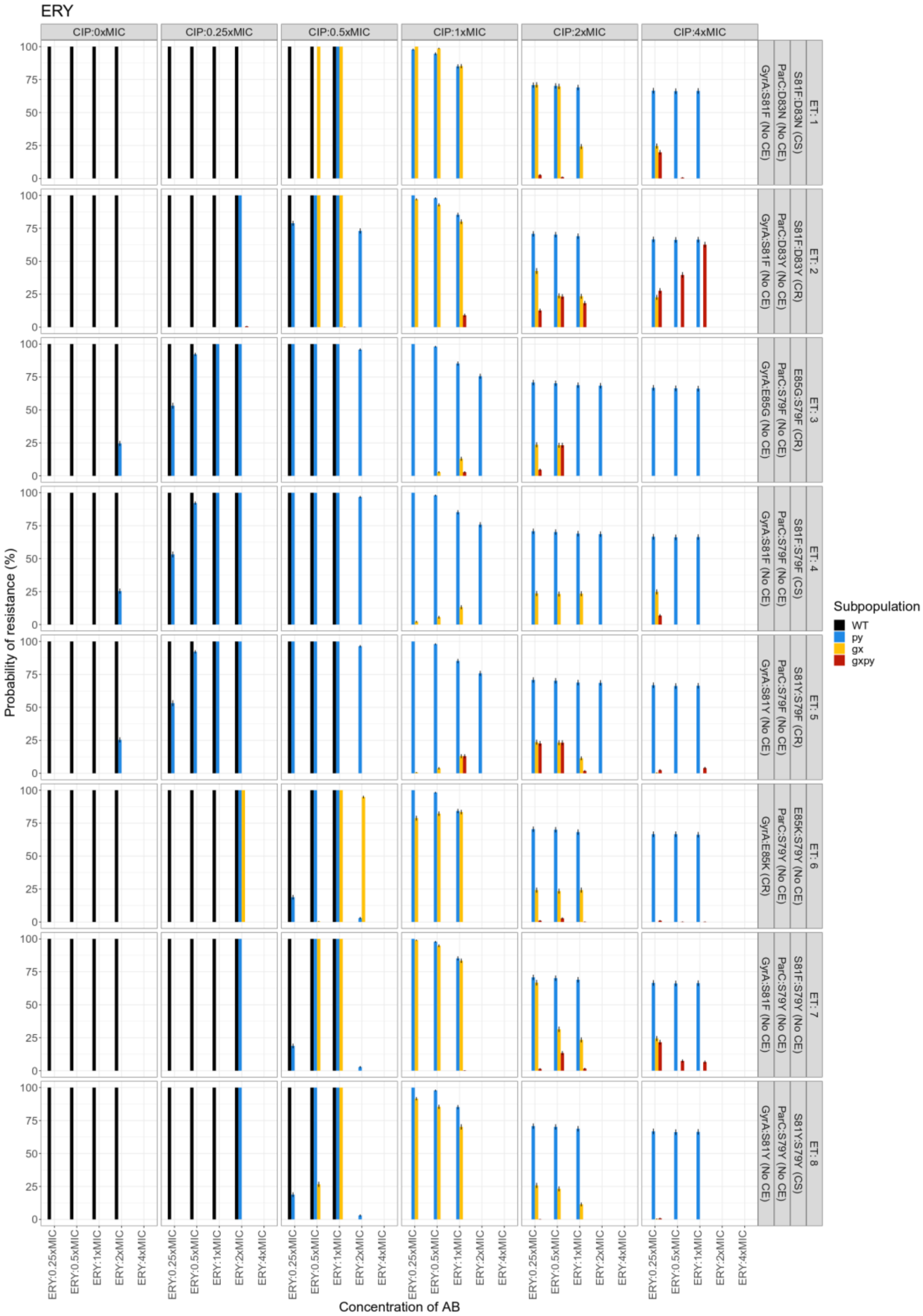
ERY
CIP:0xMIC
CIP:0.25xMIC
CIP:0.5xMIC
CIP:1xMIC
CIP:2xMIC
CIP:4xMIC
ET: 1
S81F:D83N (CS)
ParC:D83N (No CE)
GyrA:S81F (No CE)
ET: 2
S81F:D83Y (CR)
ParC:D83Y (No CE)
GyrA:S81F (No CE)
ET: 3
E85G:S79F (CR)
ParC:S79F (No CE)
GyrA:E85G (No CE)
ET: 4
S81F:S79F (CS)
ParC:S79F (No CE)
GyrA:S81F (No CE)
ET: 5
S81Y:S79F (CR)
ParC:S79F (No CE)
GyrA:S81Y (No CE)
ET: 6
E85K:S79Y (No CE)
ParC:S79Y (No CE)
GyrA:E85K (CR)
ET: 7
S81F:S79Y (No CE)
ParC:S79Y (No CE)
GyrA:S81F (No CE)
ET: 8
S81Y:S79Y (CS)
ParC:S79Y (No CE)
GyrA:S81Y (No CE)
Probability of resistance (%)
Concentration of AB
ERY:0.25xMIC
ERY:0.5xMIC
ERY:1xMIC
ERY:2xMIC
ERY:4xMIC
Subpopulation
WT
py
gx
gxpy

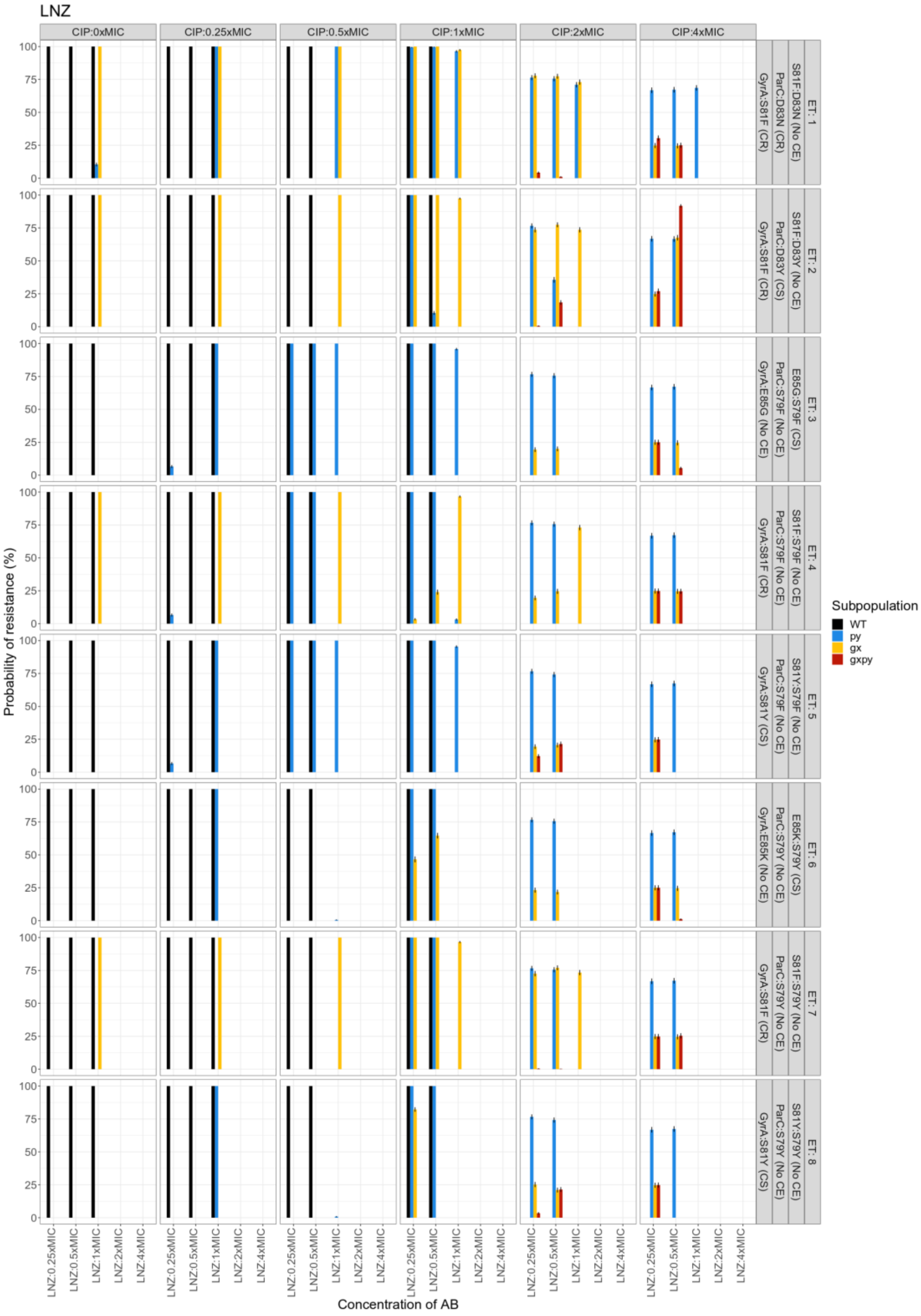

LNZ
CIP:0xMIC
CIP:0.25xMIC
CIP:0.5xMIC
CIP:1xMIC
CIP:2xMIC
CIP:4xMIC
ET: 1
S81F:D83N (No CE)
ParC:D83N (CR)
GyrA:S81F (CR)
ET: 2
S81F:D83Y (No CE)
ParC:D83Y (CS)
GyrA:S81F (CR)
ET: 3
E85G:S79F (CS)
ParC:S79F (No CE)
GyrA:E85G (No CE)
ET: 4
S81F:S79F (No CE)
ParC:S79F (No CE)
GyrA:S81F (CR)
ET: 5
S81Y:S79F (No CE)
ParC:S79F (No CE)
GyrA:S81Y (CS)
ET: 6
E85K:S79Y (CS)
ParC:S79Y (No CE)
GyrA:E85K (No CE)
ET: 7
S81F:S79Y (No CE)
ParC:S79Y (No CE)
GyrA:S81F (CR)
ET: 8
S81Y:S79Y (No CE)
ParC:S79Y (No CE)
GyrA:S81Y (CS)
Probability of resistance (%)
0
25
50
75
100
LNZ:0.25xMIC
LNZ:0.5xMIC
LNZ:1xMIC
LNZ:2xMIC
LNZ:4xMIC
Concentration of AB
Subpopulation
WT
py
gx
gxpy

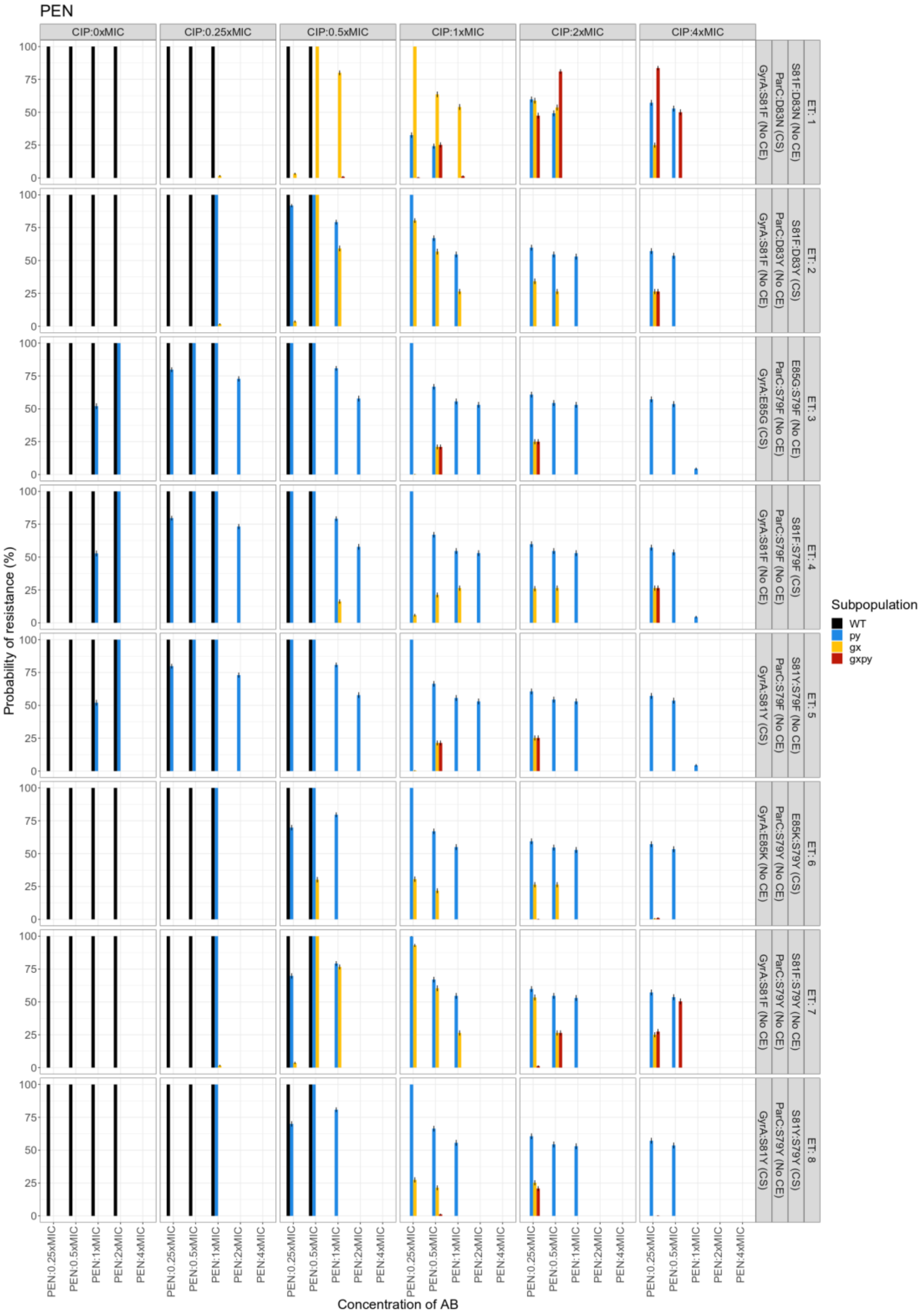


**Figure S6.** Overall treatment outcomes of simulated ciprofloxacin mono- or combination with erythromycin (ERY), linezolid (LNZ), or penicillin (PEN).

Simulation outcomes treated with CIP in combination with ERY, LNZ, or PEN with average unbound steady state concentrations ($C_{ss}$) around the MIC of the WT population ($C_{ss}$ 0.25 - 4 x MIC). Mutant subpopulations included a *gyrA* mutant gx, a *parC* mutant py, and the double-allele mutant gxpy. The treatments were evaluated on eight different trajectories (ET) leading to high-level FQ-resistance. End-of-treatment probability of resistance (mutants) and treatment failure (WT), were calculated for the different treatments (panels), evolutionary trajectories (rows), and subpopulations (indicated by color) separately. Collateral effects (CE) are indicated per mutant in corresponding row panel, where CS is collateral sensitivity and CR collateral resistance, if applicable.

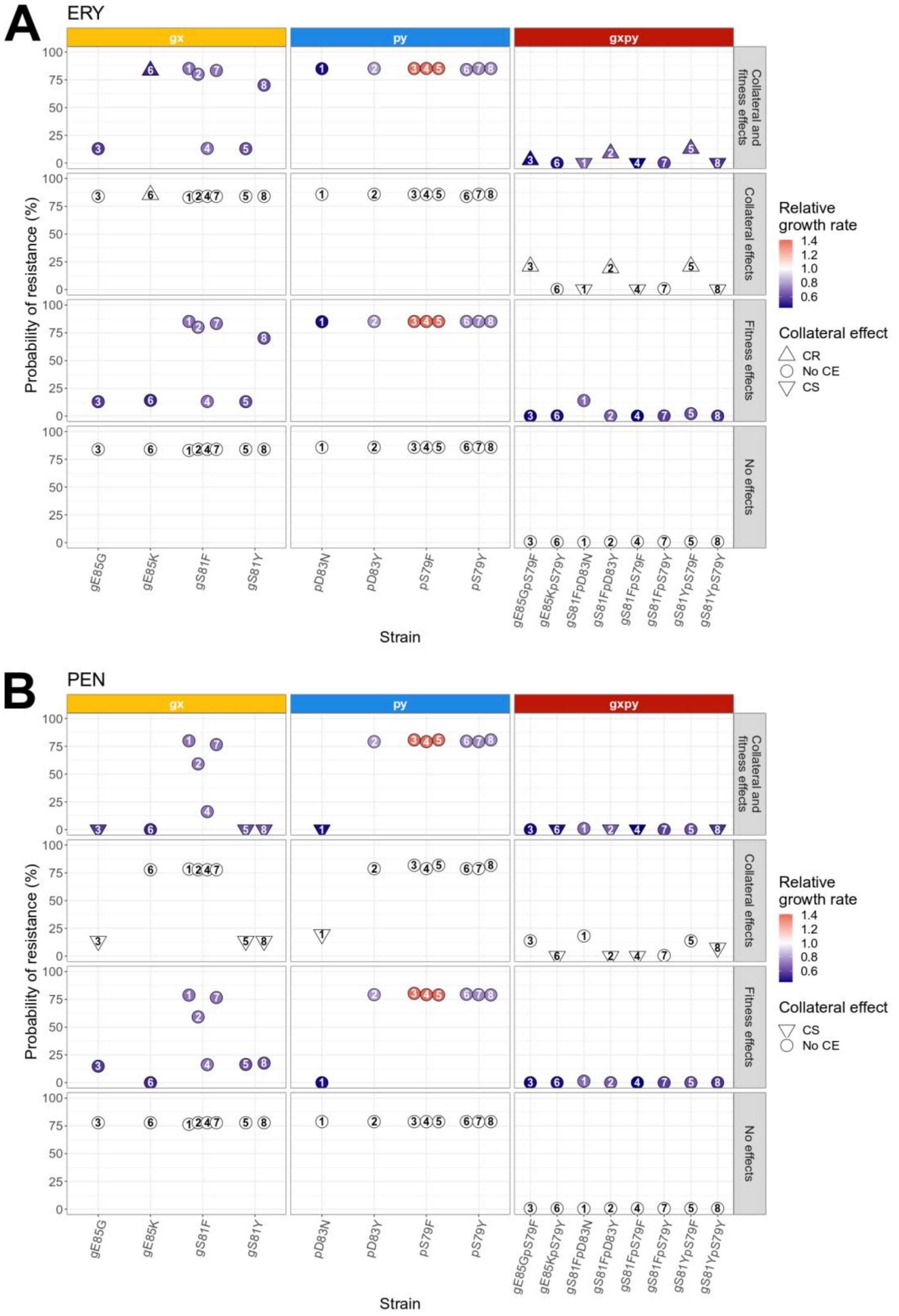


**Figure S7.** Relationship between collateral effect, relative growth rate, and probability of resistance under antibiotic combinations. Simulation of the eight defined *S.*

*pneumoniae* evolutionary trajectories (indicated by number) treated with a combination ciprofloxacin (CIP) and **A**: erythromycin (ERY) or **B:** penicillin (PEN). Each unique strain is represented on the x-axis. The collateral effect (CE) of the second drug is depicted using shapes, where tringle point up is collateral resistance (CR), pointing down is collateral sensitivity, and circle represents no CE. The probability of resistance at the end of treatment is shown on the y-axis. Colors indicate the relative growth rate of each mutant compared to the WT. Each panel column denotes a FQ-resistant subpopulation (gx, py or gxpy) with the top panels representing simulations were both collateral or fitness effects were included, top mid panels based solely on the experimentally determined collateral effects, lower mid panels based solely on the experimentally determined relative growth rate, and the lower panel simulations where no collateral and fitness effects were included. The steady state concentrations (Css) used were 1 x MIC CIP + 1 x MIC ERY or 0.5 x MIC CIP + 1 x MIC LNZ, respectively.

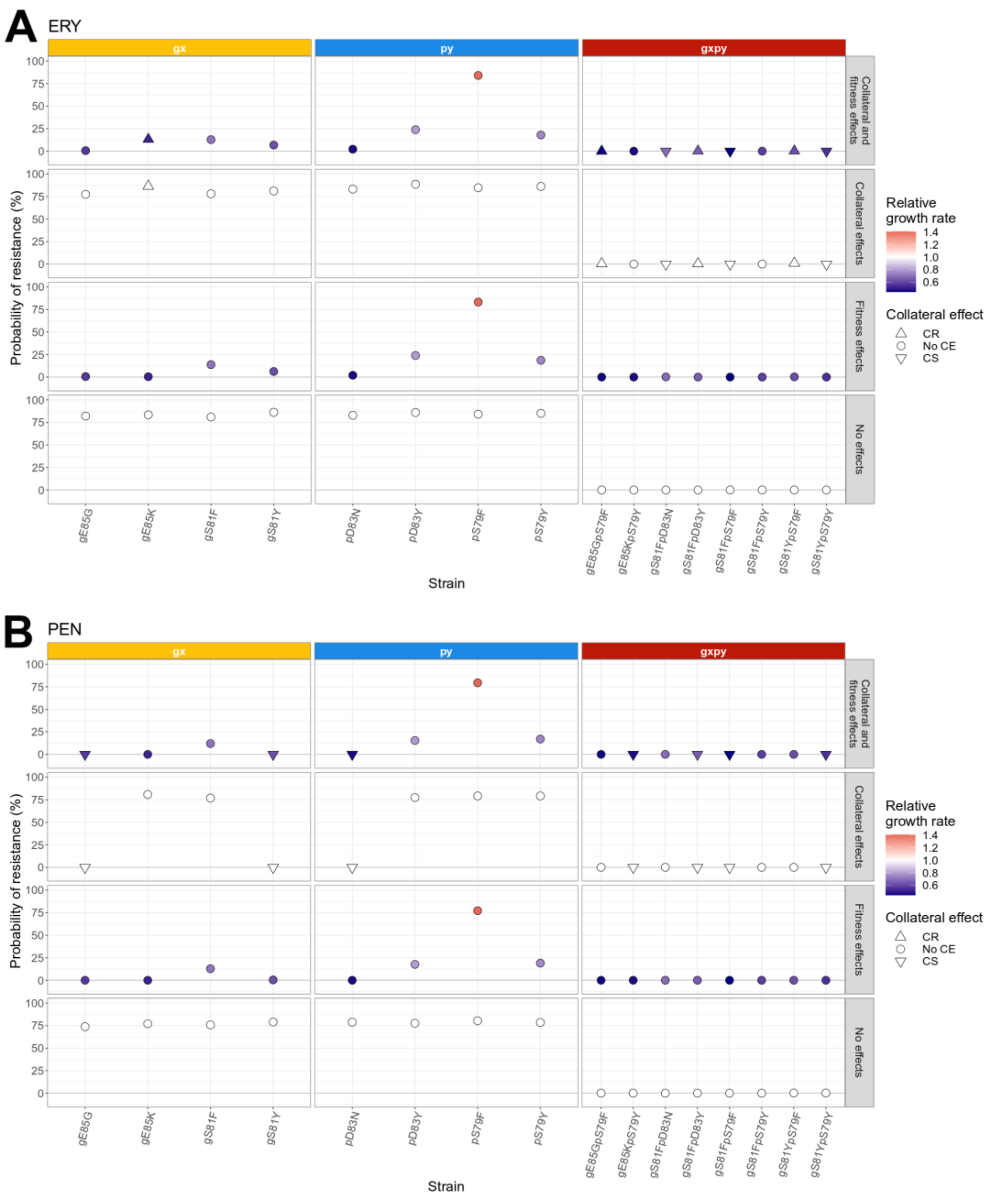


**Figure S8.** Relationship between collateral effect, relative growth rate, and probability of resistance for specific mutants under antibiotic combinations. Simulation of resistance evolution in *S. pneumoniae* treated with a combination ciprofloxacin (CIP) and erythromycin (ERY; A) or penicillin (PEN; B). These free-for-all simulations allows for the simultaneous evolution of 16 different FQ-resistant subpopulations (four gx, four py and eigth gxpy). Each unique strain is represented on the x-axis. The collateral effect (CE) of the second drug is depicted using shapes, where tringle point up is collateral resistance (CR), pointing down is collateral sensitivity, and circle represents no CE. The probability of resistance at the end of treatment is shown on the y-axis. Colors indicate the relative growth rate of each mutant compared to the WT. Each panel column denotes a FQ-resistant mutant subpopulation (gx, py or gxpy) with the top panels representing simulations were both collateral or fitness effects were included, top mid panels based solely on the experimentally determined collateral effects, lower mid panels based solely on the experimentally determined relative growth rate, and the lower panel based where no collateral and fitness effects were included. The steady state concentrations (Css) used were 1 x MIC CIP + 1 x MIC ERY or 0.5 x MIC CIP + 1 x MIC LNZ, respectively.

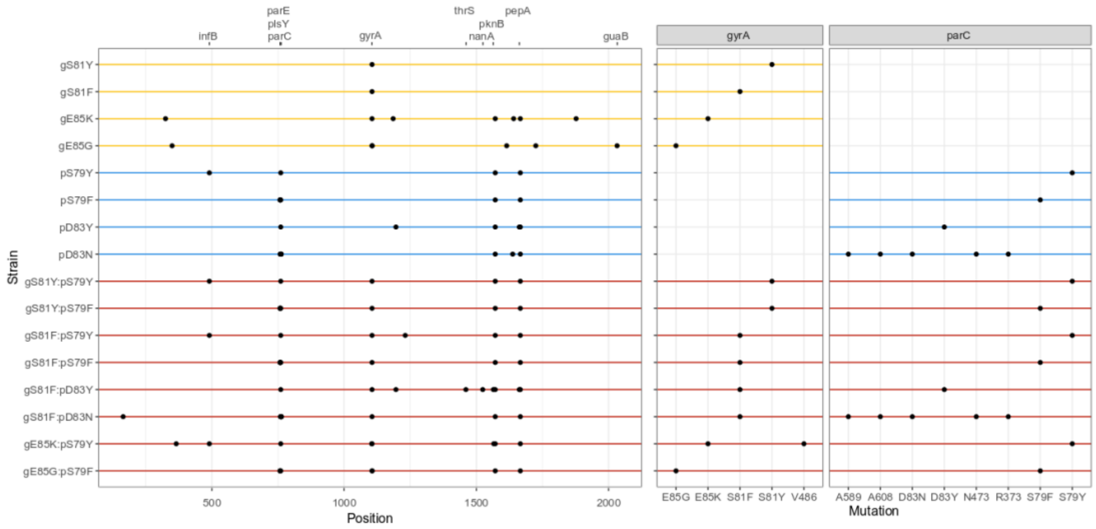


**Figure S9.** SNP calling for each generated FQ-resistance strain comparing to the WT strain. Color indicates *gyrA* mutants (yellow), *parC* mutants (light blue), and their respective double-allele mutant (red).

### Appendix 1.

#### Derived concentration-effect relationships

We digitized early phase data (≤ 6 h after start of experiment) from previously performed *in vitro* static time-kill experiments treating *S. pneumoniae* with ciprofloxacin, erythromycin, linezolid, or penicillin. Summary of the collected data can be found in **Table A1.1**. The data was subsequently used for fitting antibiotic specific pharmacodynamic models. All fitted models could satisfactorily describe the observed data (**Figure A1.1**). Penicillin was estimated to have the largest maximal drug effect (Gmin = - 1.28) and linezolid the lowest (Gmin = - 0.371), while antibiotic differences were seen for the Hill factor ranged from 0.609 (ERY) to 2.81 (LNZ) (**Table A1.2**, **Figure A1.2**).

**Table A1.1.** Overview of collected litterateur data of *in vitro* time-kill studies on *S. pneumoniae*.

| Drug | Concentration range | N conc | MIC | ID | Reference |
|---|---|---|---|---|---|
| CIP | 0.125-16 µg/mL | 8 | 2.0 µg/mL | 1 | (29) |
| CIP | 0.25-32 µg/mL | 8 | 4.0 µg/mL | 2 | (29) |
| CIP | 0.25-32 µg/mL | 8 | 4.0 µg/mL | 3 | (29) |
| CIP | 0.125-16.0 µg/mL | 8 | 2.0 µg/mL | 4 | (30) |
| CIP | 0.002-0.064 µg/mL | 5 | 0.008 µg/mL | 5 | (31) |
| ERY | 0.004-0.5 µg/mL | 8 | 0.06 µg/mL | 1 | (29) |
| ERY | 2-256 µg/mL | 8 | 32 µg/mL | 2 | (29) |
| ERY | 0.5-64 µg/mL | 8 | 8.0 µg/mL | 3 | (29) |
| ERY | 0.05-0.4 µg/mL | 3 | 0.1 µg/mL | 4 | (32) |
| LNZ | 1.0-20 µg/mL | 4 | 1.0 µg/mL | 1 | (33) |
| PEN | 0.001-0.125 µg/mL | 9 | 0.015 µg/mL | 1 | (29) |
| PEN | 0.015-2.0 µg/mL | 9 | 0.25 µg/mL | 2 | (29) |
| PEN | 0.125-16.0 µg/mL | 9 | 2.0 µg/mL | 3 | (29) |
| PEN | 0.250-32.0 µg/mL | 9 | 4.0 µg/mL | 4 | (30) |
| PEN | 4 – 32 µg/mL | 4 | 8.0 µg/mL | 5 | (34) |
| PEN | 0.008-0.064 µg/mL | 4 | 0.016 µg/mL | 6 | (34) |

**Table A1.2.** Pharmacodynamic model parameters.

| **Drug** | **Gmin** (95%CI) [RSE%] | **Hill factor** (95%CI) [RSE%] |
|---|---|---|
| Ciprofloxacin | 1.17 (1.12, 1.22) [14.9] | 1.55 (1.36, 1.75) [14.8] |
| Erythromycin | 1.02 (0.51, 2.05) [1680] | 0.636 (0.421, 0.96) [46.5] |
| Linezolid | 0.371 (0.33, 0.418) [6.09] | 2.99 (0.041-217) [159] |
| Penicillin | 1.28 (1.15, 1.43) [22.2] | 1.87 (1.58, 2.21) [13.8] |

CI confidents interval, RSE relative standard error

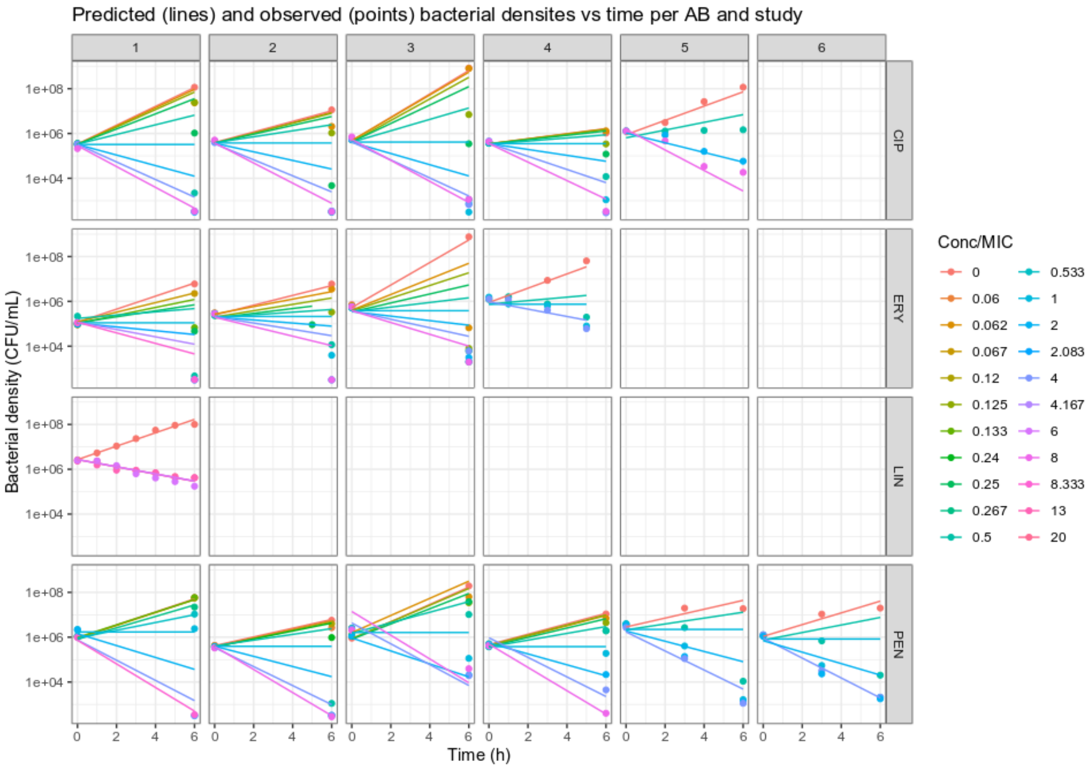


**Figure A1.1.** Predicted (lines) and observed (points) bacterial densities of *S. pneumoniae* treated with ciprofloxacin (CIP), erythromycin (ERY), linezolid (LIN), penicillin (PEN), where color indicate antibiotic concentration normalized over the MIC.

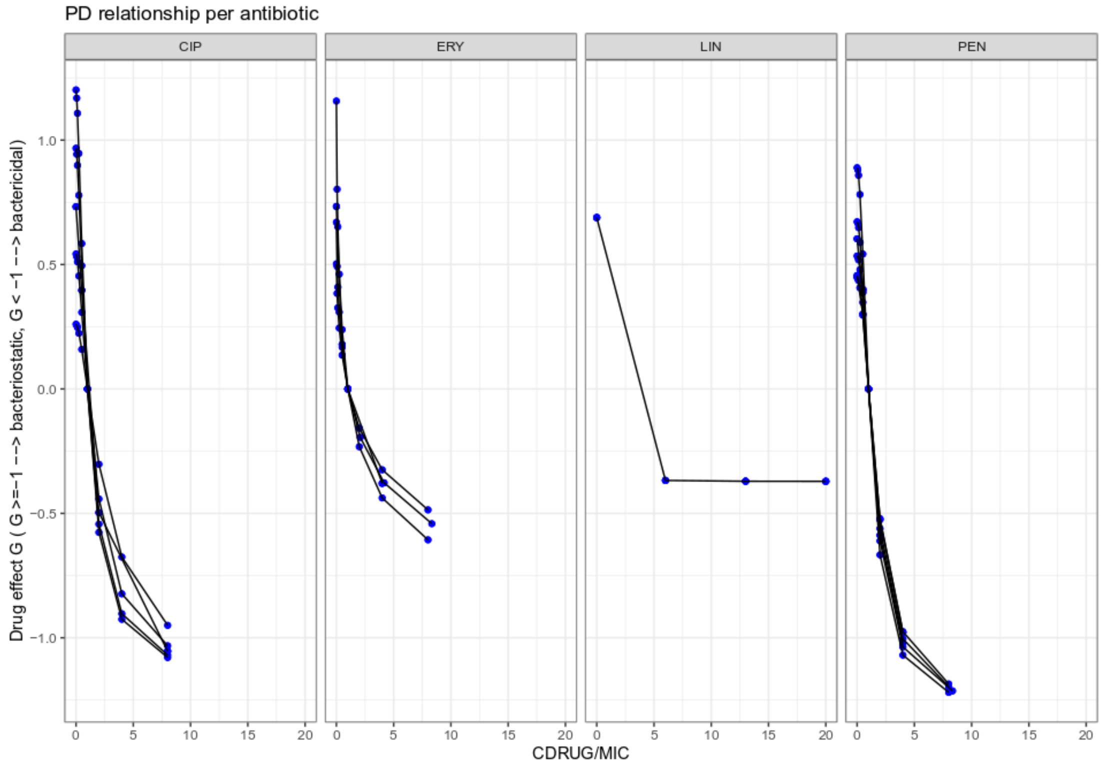


**Figure A1.2.** Pharmacodynamic concentration-effect-relationship of ciprofloxacin (CIP), erythromycin (ERY), linezolid (LIN), penicillin (PEN).

**Appendix 2.**

To assess the interaction of ciprofloxacin with linezolid, erythromycin, and penicillin, respectively, we performed minimal time-kill assays. Subsequently, we used the generated data and fitted a non-linear model (Figure A2.1) form which we obtain the interaction terms summarized in Table A2.1.

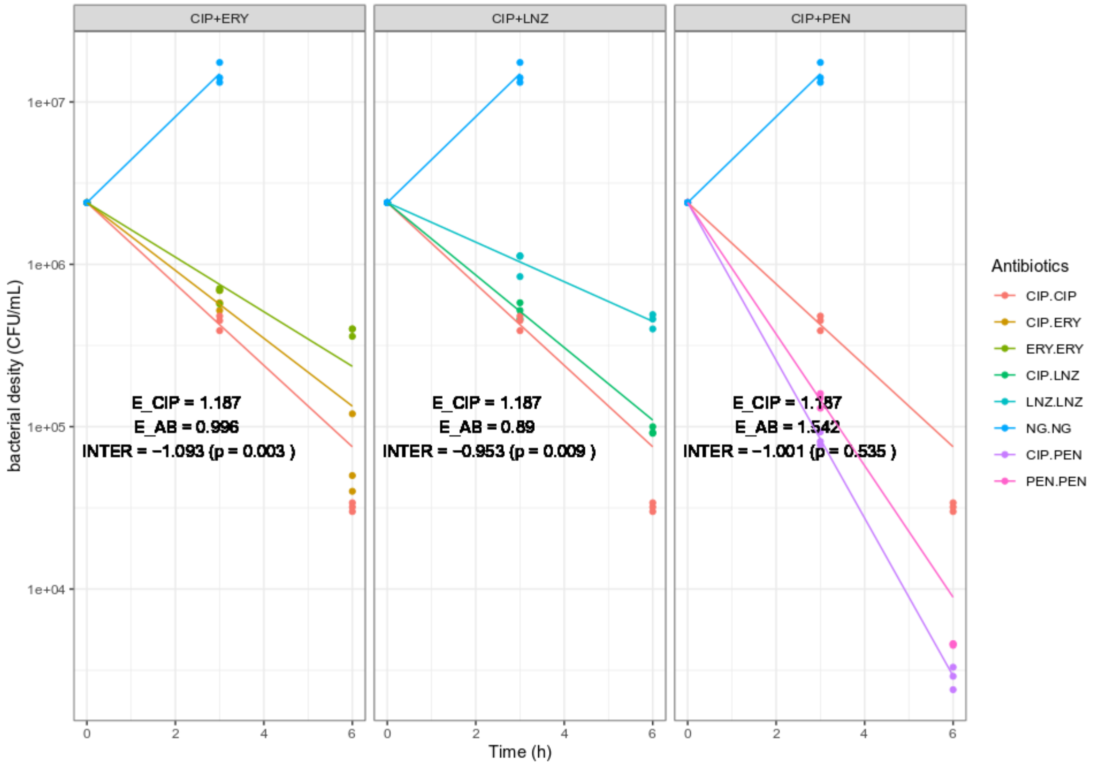


Figure A2.1. Observed (points) and predicted (lines) bacterial densities of *Streptococcus pneumoniae* D39 treated with ciprofloxacin (CIP), linezolid (LNZ), penicillin (PEN), or erythromycin (ERY).

**Table A2.1** Derived interaction parameters

| Antibiotic | Effect | Interaction effect | Relative interaction term | p |
|---|---|---|---|---|
| CIP | 1.19 | NA | NA | NA |
| LNZ | 0.89 | -0.95 | -0.46 | 0.0087 |
| PEN | 1.54 | -1.00 | -0.37 | 0.5350 |
| ERY | 1.00 | -1.09 | -0.50 | 0.0033 |

CIP = ciprofloxacin, LNZ = linezolid, PEN = penicillin, ERY = erythromycin
